# Identifying associations in dense connectomes using structured kernel principal component regression

**DOI:** 10.1101/242982

**Authors:** Weikang Gong, Fan Cheng, Edmund T. Rolls, Lingli Zhang, Stefan Grünewald, Jianfeng Feng

## Abstract

A powerful and computationally efficient multivariate approach is proposed here, called structured kernel principal component regression (sKPCR), for the identification of associations in the voxel-level dense connectome. The method can identify *voxel*-phenotype associations based on the voxels’ whole-brain connectivity pattern, which is applicable to detect linear and non-linear signals for both volume-based and surface-based functional magnetic resonance imaging (fMRI) data. For each voxel, our approach first extracts signals from the spatially smoothed connectivities by structured kernel principal component analysis, and then tests the voxel-phenotype associations via a general linear model. The method derives its power by appropriately modelling the spatial structure of the data. Simulations based on dense connectome data have shown that our method can accurately control the false-positive rate, and it is more powerful than many state-of-the-art approaches, such as the connectivity-wise general linear model (GLM) approach, multivariate distance matrix regression (MDMR), adaptive sum of powered score (aSPU) test, and least-square kernel machine (LSKM). To demonstrate the utility of our approach in real data analysis, we apply these methods to identify voxel-wise difference between schizophrenic patients and healthy controls in two independent resting-state fMRI datasets. The findings of our approach have a better between-sites reproducibility, and a larger proportion of overlap with existing schizophrenia findings. Code for our approach can be downloaded from https://github.com/weikanggong/vBWAS.

## 1 Introduction

Functional connectivity analysis using resting-state functional magnetic resonance imaging (fMRI) data has become increasingly popular in the last few years (e.g. [Smith et al., 2015; Finn et al., 2015]), and the advances have led to many investigations of functional dysconnectivity between brain areas in neurodegenerative and psychiatric brain diseases [Gong and He, 2015; Romme et al., 2017]. Voxelbased functional connectivity analysis has also emerged [Cheng et al., 2015b, 2016; Satterthwaite et al., 2015; Kaczkurkin et al., 2017]. However, designing methods for an exploration of associations between the whole-brain voxel-level connectome and phenotypes is a challenging task, and well-developed approaches are usually designed for parcellation-based or seed-based connectivity studies [Meskaldji et al., 2013; Bellec et al., 2015; Xia and He, 2017].

The most popular method for functional connectivity analysis is the massive univariate general linear model (GLM) approach. It uses a general linear model to test the association between each voxel-voxel connectivity and the phenotype of interest, and then corrects for multiple comparison by methods, such as Bonferroni correction or false-discovery rate [Benjamini and Hochberg, 1995], to locate the significant signals. The major advantage of this approach is that it can provide us the exact location of the signals. However, the large number of hypothesis tests requires a stringent multiple correction threshold, which usually decreases the power and increases the potential for false-positive discoveries. In addition, univariate approaches only test the linear marginal association between connectivities and phenotypes. Important higher-order information, such as the co-contribution of a set of functional connectivities and the non-linear associations, is usually ignored by this method.

In recent years, many improvements over univariate method have been proposed. These approaches usually adopt a global association test to achieve higher power. In other words, they test whether the signal is present somewhere in a set of functional connectivities rather than localizing it. For example, the network based statistic (NBS) and spatial pairwise clustering (SPC) [Zalesky et al., 2010, 2012] are two popular approaches. They first perform statistical tests on each connectivity, and then use a permutation-based method to test whether the size of the suprathreshold connectivities is larger than by chance. The above two approaches are based on the assumption that the signals form a connected graph in the connectome. Pan et al. [2014]; Kim et al. [2014, 2015] proposed the adaptive sum of powered score (aSPU) test and its extensions. This approach first assigns a score to measure the association between a phenotype and an individual connection. It then combines the individual scores into a summary statistic and uses a permutation test to access the significance. The multivariate distance matrix regression [Shehzad et al., 2014] is an ANOVA-like nonparametric multivariate approach, which directly tests the association between a phenotype of interest and a between-subject distance matrix estimated using the functional connectivity data. Other approaches include [Simpson and Laurienti, 2015; Chen et al., 2015; Fiecas et al., 2017; Meskaldji et al., 2015; Belilovsky et al., 2016]. However, in the context of voxel-level connectivity analysis, the above approaches have three major drawbacks: First, the spatial structure of the dense connectome, which is structurally and smoothly correlated, is different from the region-level connectome. Therefore, as shown in our analysis, the un-modelled spatial noise usually decreases their power. Second, most of these approaches use non-parametric permutation to get voxel/connectivity-wise p-values, which is computationally expensive, as the dimensionality of the feature space grows quadratically with the number of voxels (*O*(*p*^2^)) and the computation time grows quadratically with the sample size (*O*(*n*^2^)). In addition, a huge number of permutations should be performed to get a reliable estimation of a small p-value (e.g. more than 10^5^ permutations are required to get a p-value of 10^-5^). Third, these approaches can only detect linear association signals. Important non-linear signals may be missed by them.

In this paper, a new multivariate approach is proposed to overcome the above problems. It is designed specifically for dense connectome and applicable for both volume-based and surface-based fMRI data. Our approach evaluates, for each voxel, the simultaneous contribution of its whole-brain connectivities to a phenotype of interest. It has three steps: (1) Extracting important features from the data using a newly-developed structured kernel principal component analysis (sKPCA) approach; (2) Testing the association between the low-dimensional features (principal components) and the phenotype of interest using general linear regression; (3) Controlling the voxel-wise family-wise error rate (FWER) using an efficient non-parametric permutation procedure. The sKPCA is an extension of the widely-used principal component analysis (PCA) [Jolliffe, 2002] method. Unlike the PCA method assuming independent and identically distributed noise structure, sKPCA assumes a spatially correlated noise structure among functional connectivities. We will show in our analysis that, compared with other approaches, sKPCR is the most powerful one by utilizing the spatial information. Moreover, a non-linear extension is also developed based on the idea of kernel principal component analysis [Schölkopf et al., 1997]. In addition, sKPCR is highly efficient, thus, it can handle huge datasets with high-resolution data while other approaches can not. Other attractive features of our approach include: 1) applicability for both categorical (e.g. disease status) and continuous variables (e.g. IQ, symptom score), 2) covariate effects (e.g. age, gender, motion) can be considered, 3) the parameters of the model are easily specified (e.g. the number of general principal components and the covariance structure of the dense connectome).

The remainder of the paper is organized as follows. We first describe the details of our method (see Figure 1 for a graphical overview). We then conduct comprehensive simulations to compare the proposed approach with several state-of-the-art methods discussed above, including their power, false-positive rate and computation time. Finally, we evaluate and compare the performance with real data by applying it to identify voxel-wise differences between schizophrenic patients and healthy controls, and test the association between schizophrenic patients and the Positive and Negative Syndrome Scale Score (PANSS) [Kay et al., 1987]. The code for our approach can be downloaded from https://github.com/weikanggong/vBWAS.

**Figure 1:**
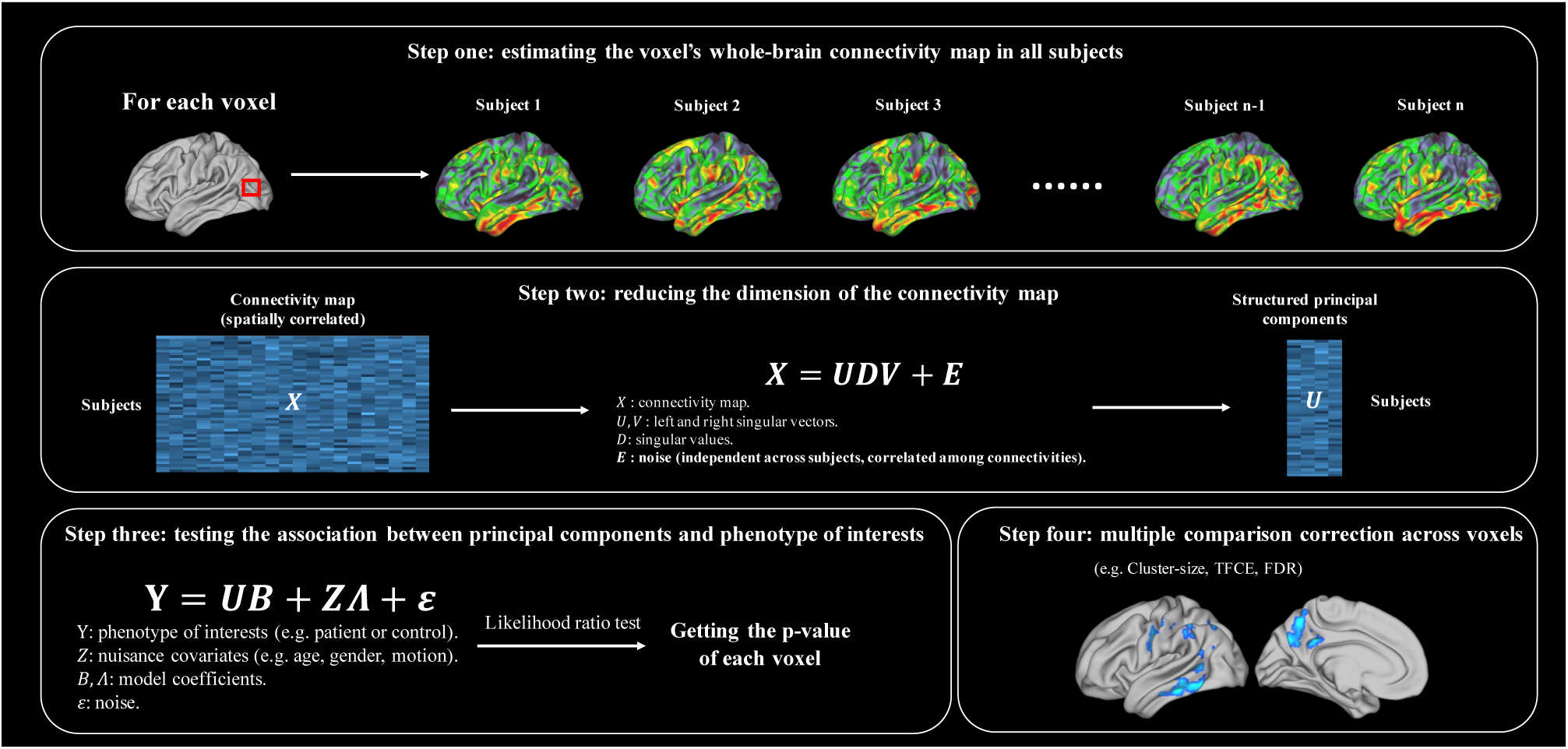
An overview of the structured kernel principal component regression in dense-connectome association study. First, for each voxel and each subject, the whole-brain functional connectivity map is computed. Second, a newly-developed dimension reduction technique is applied to extract important features in this connectivity map. Then, a general linear model is fitted to test the association between a phenotype of interest and this voxel. Finally, voxel-wise multiple correction is performed to identified significant clusters.

## 2 Method

### 2.1 Structured kernel principal component analysis

Principal component analysis [Jolliffe, 2002] (PCA) is one of the most popular dimension reduction and feature extraction approaches. Given a row-wise Gaussian normalized data matrix *X* ∈ ℝ*^n^*^×^*^p^*, where *n* is the number of samples and *p* is the number of features (*p* ≫ *n*), the probabilistic model of PCA is to find the best linear rank-*k* approximation of the original data X:

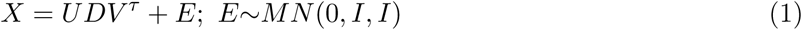

where *U* ∈ ℝ*^n^*^×^*^k^*, *V* ∈ ℝ*^p^*^×^*^k^* are matrices of the left and right singular vectors of *X*, *D* ∈ ℝ*^k^*^×^*^k^* is a diagonal matrix of the singular values of *X*, and *E* is the noise matrix which is subject to a zero-mean matrix normal distribution *MN* (0, *I*, *I*), i.e., the noise is independent between subjects and features. There are two major drawbacks of the conventional PCA when applied to analyse high dimensional neuroimaging data in our analysis: (1) the PCA assumes independent and identically distributed (iid) noise for both the rows and columns of the X. However, the assumption may break down because most features (e.g. voxels, functional connectivity) are spatially correlated; (2) the PCA can only perform linear dimension reduction and feature extraction, and many important non-linear factors may be missed by this method. Therefore, we propose a structured kernel principal component analysis (sKPCA) approach. It can model the spatial dependence between noise terms and extract both linear and non-linear features.

To address the first problem, we allow the noise terms between features to be dependent with each other, thus, we propose the following modification of the PCA model (1), which we call structured principal component analysis (sPCA), as:

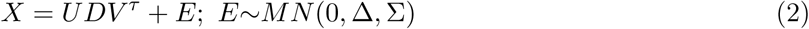

where the noise matrix *E* is subject to a zero-mean matrix normal distribution *MN* (0, ∆, Σ), the subject-wise covariance matrix is ∆ ∈ ℝ*^n^*^×^*^n^*, and the feature-wise covariance matrix is ∆ ∈ ℝ*^p^*^×^*^p^*. In this model, we allow noises among features and samples to be correlated, but in the subsequent applications, we assume that the samples are independent, i.e., ∆ = *I*.

It is well-known that the PCA problem (1) can be solved by minimizing the reconstruction error with respect to the Frobenius norm: minimize 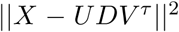, subject to *U^τ^U* = *I*, *V^τ^V* = *I* and 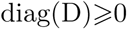. Similar to the PCA model (1), the sPCA model (2) is equivalent to solve the following optimization problem with respect to *U*, *D* and *V*:

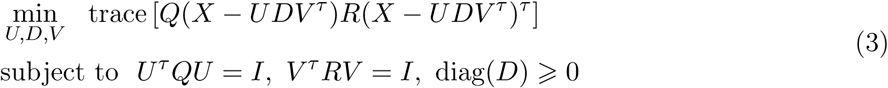

where *Q* = ∆^-1^ and *R* = Σ^-1^. Escoufier [1977]; Allen et al. [2014] have shown that, if we let 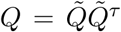 and 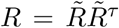, then the optimization problem (3) can be solved by performing singular value decomposition on 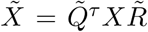. Let 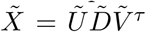, the solution is 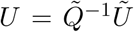, 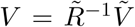 and 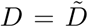. In our case, let *Q* = *I*, the *U* = (*u*_1_,…, *u_k_*) can be obtained by performing an eigenvalue decomposition on matrix 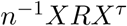. This is known as a generalization of the maximizing the projected variance formulation of PCA. By sorting the eigenvalue λ in a descending order, the *k*-th principal component *u_k_* is the *k*-th eigenvector of *n*^-1^*X^τ^RX*. The variance explained by the *k*-th principal component is *λ_k_*/trace(*n*^-1^*X^τ^RX*). In (2) or (3), the *R* do not need to be estimated from the data, but to be pre-specified based on the topological structure of the data. We will introduce how to specify the *R* in the next Section.

The sPCA can be further generalized to perform either linear or non-linear PCA by using the kernel tricks. Let 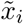 be the *i*-th ‘weighted sample’, i.e., the *i*-th row of 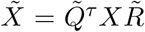, we first perform a non-linear mapping of the sample 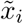 to the high dimensional feature space as 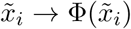. Now, we assume that each 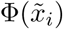 has been mean centered in the feature space and we will return to this point later. Motivated by the model (3), we perform a PCA in the mapped high-dimensional feature space by maximizing the projected variance as:

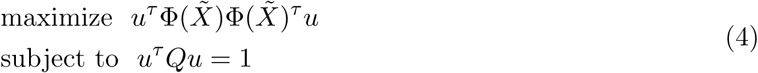

Similar to the kernel principal component analysis [Schölkopf et al., 1997], the optimization problem (4) can be solved by first performing a mean normalization of the kernel matrix *K* ∈ ℝ*^n^*^×^*^n^*, where *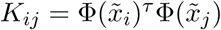*, by:

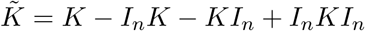

where *I_n_* is an *n* × *n* matrix with each elements takes the value 1/*n*. Then, we solve the eigenvalue problem:

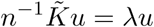

and obtains *n* eigenvalues in a descending order as (λ_1_,…,*λ_n_*) and the corresponding eigenvectors (*u*_1_,…, u_n_). The *k*-th principal component is the *k*-th eigenvector *u_k_*. The variance explained by the *k*-th principal component is given by 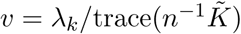.

Similar to most other kernel-based approaches, all the computations can be expressed in terms of the kernel matrix. When using the linear kernel, the sKPCA is exactly the same as sPCA. In addition, for many commonly used kernels, we even do not need to estimate *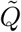* and *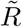.* For example, we can calculate *X*\* = *QXRX^τ^Q,* and the polynomial kernel can be calculated as 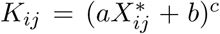, the sigmoid kernel can be calculated as 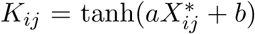, and the gaussian kernel can be calculated as 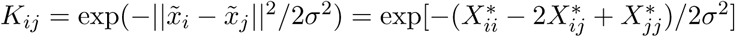.

### 2.2 The choice of sKPCA parameters

There are many developed methods to determine the number of principal components for conventional PCA, such as the ratio estimator [Lam and Yao, 2012; Li et al., 2017b], the information criteria approaches [Bai and Ng, 2002,2007], the distribution-based approach [Choi et al., 2014] or just by the amount of variance explained (e.g. 80%) or the average variance explained. These methods can be easily extended to the sKPCA framework. In the context of connectivity analysis, we propose to use the following method, which usually achieves high power in subsequent association studies empirically. The number of principal components selected is:

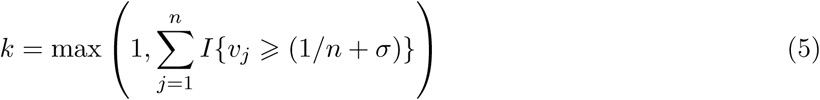

where *v_j_* is the proportion of explained variance of the *j*-th principal components, 1/*n* is the average proportion of explained variance of *n* principal components, *σ* is the standard deviation of the proportion of explained variances of *n* principal components, and *I*{·} is the indicator function. This means that we select the principal components whose explained variances are larger than one standard deviation of the mean variance explained.

There are also many possible choices for the covariance matrix [Allen et al., 2014; Ramsay, 2006], and we mainly introduce three of them in this paper. The first one is the Graph Laplacian operator, which has been widely used in Bayesian task-activation studies (e.g. [Penny et al., 2005; Flandin and Penny, 2007; Sidén et al., 2017]). It is also known as the inverse covariance operator [Allen et al., 2014; Ramsay, 2006]. To define the Graph Laplacian operator *G*, we first define the feature-feature adjacency matrix *A* as a binary matrix such that *a_ij_*· = 1 if the spatial distance between feature *i* and *j* (*i* ≠ *j*) equals one (or feature *i* and *j* are spatial neighbours) and *a_ij_*· = 0 otherwise. Based on *A*, we can define G as, for feature *i* and *j,* 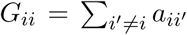 and *G_ij_* = − *a_ij_* if *i* ≠ *j*. The second one is the normalized Graph Laplacian operator *G*^⋆^. Based on *A*, it is defined as 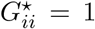 and 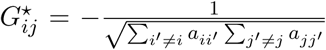 if *i* = *j*. The third one is called the Gaussian random field operator. It assumes that the noise covariance between two features is a functional of their spatial distance: 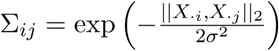 where 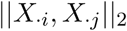 represents the spatial Euclidian distance between feature *i* and features *j* in the volume space, and 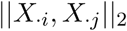 can also be the geodesic distance in surface space. The *σ* can be specified based on the estimated Full width at half maximum (FWHM) of the images using the relationship 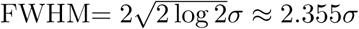.

In the subsequent real data analysis, we will show that sKPCR is very stable for selecting different numbers of principal components and covariance operators.

### 2.3 Identifying connectome-wide associations

We extend sKPCA to structured kernel principal component regression (sKPCR) for identifying the connectome-wide associations. In our study, the individual-level brain functional network is estimated by the Fisher’s Z transformed Pearson correlation coefficient between every pair of voxels’ BOLD signal time series. Let *n* be the number of subjects in a study, and *p* be the number of voxels, thus, there are a total number of *p*(*p* − 1)/2 functional connectivities in each individual’s brain network and *p* − 1 functional connectivities connecting a voxel to all other voxels across the whole brain. Let *Y* ∈ ℝ*^n^*^×1^ be the phenotype of interest of *n* subjects (e.g. disease status, clinical symptoms) and *Z* ∈ ℝ*^n^*^×^*^q^* be the nuisance covariates (e.g. age, gender, motions). Our aim is to test, for each voxel, whether the phenotype of interest *Y* is associated with the voxel’s whole-brain functional connectivity pattern *X* ∈ ℝ*^n^*^×(^*^p^*^-1)^, conditioned on the nuisance covariates *Z*. Since connectivity is of ultra-high dimensionality (e.g. for each voxel, there are 10^4^ to 10^5^ whole-brain functional connectivities, but only a few hundred samples), the basic idea of our model is to first extract important low-dimensional features (principal components) in the data by sKPCA, and then test the association between the extracted principal components and the phenotype of interest.

For each voxel, our method proceeds with two steps: (1) Performing sKPCA on *X* and extracting the top k principal components *U* = (*u*_1_*,…,u_k_*); (2) Fitting a general linear model to test the association between the principal components *U* and the phenotype *Y* as:

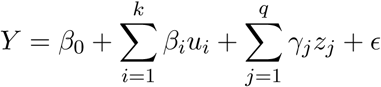

A multivariate linear model is used if *Y* is a continuous variable, and an *F* test is used to compare the full model with the null model, 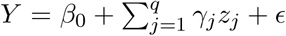. A logistic regression model is used if *Y* is a binary variable, and a likelihood ratio test is performed to compare the full model with the null model 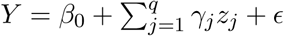, and a chi-square statistic with k degree of freedom can be obtained.

After getting the voxel-wise p-values, we can use a non-parametric permutation approach [Nichols and Holmes, 2002] or false-discovery rate method [Benjamini and Hochberg, 1995] to perform multiple comparison correction. For permutation-based approaches, we implement a fast permutation procedure for the general linear model based on [Winkler et al., 2014; Conroy and Sajda, 2012]. All popular voxel-wise inference methods can be used here, including peak-level inference [Worsley et al., 1996], cluster-size inference [Friston et al., 1994], cluster-mass inference [Zhang et al., 2009], and threshold-free cluster enhancement [Smith and Nichols, 2009].

### 2.4 Simulation study: comparison with other methods

#### 2.4.1 Data

We use two resting-state fMRI datasets to evaluate different methods: 281 subjects from the Southwest University (SWU) dataset in the International Data-sharing Initiative (IDNI, http://fcon_1000.projects.nitrc.org/indi/retro/southwestuni_qiu_index.html); (2) 150 subjects from the Human Connectomes project (HCP, https://www.humanconnectome.org/). The detailed data acquisition method is illustrated in the Appendix A. All the subjects are healthy adults with similar demographic information.

The data from SWU are preprocessed using standard volume-based fMRI pipeline. For each individual, the preprocessing steps include: slice timing correction (FSL slicetimer), motion correction (FSL mcflirt), spatial smoothing by a 3D Gaussian kernel (FWHM = 6 mm), despiking motion artifacts using BrainWavelet Toolbox [Patel et al., 2014], registering to 4 × 4 × 4 mm^3^ standard space by first aligning the functional image to the individual T1 structure image using boundary based registration (BBR, [Greve and Fischl, 2009]) and then to standard space using FSL’s linear and non-linear registration tool (FSL flirt and fnirt), regressing out nuisance covariates including 12 head motion parameters (6 head motion parameters and their corresponding temporal derivatives), white matter signal, cerebrospinal fluid signal, band-pass filtering (0.01-0.1 Hz) using AFNI (3dTproject). All the images were manually checked to ensure successful preprocessing. Finally, 14364 grey matter voxels located in each subject’s cerebrum are extracted for the subsequent analysis.

The data from HCP-S900 are preprocessed using *fMRIsurface* minimal preprocessing pipeline [Glasser et al., 2013; Smith et al., 2013]. The basic steps include: corrected for spatial distortions caused by gradient nonlinearity; corrected for head motion by registration to the single band reference image; corrected for *B*_0_ distortion; registered to the T1w structural image; and the global intensity were normalised. Then, independent component analysis (ICA) was run using MELODIC with automatic dimensionality estimation [Beckmann and Smith, 2004]. These components are fed into FIX [Salimi-Khorshidi et al., 2014], which classifies components into ‘good’ vs. ‘bad’. Bad components are removed from the data. From this resulting volume time-series, the data were mapped onto the standard 32k Conte69 cortical surface using Multimodal Surface Matching approach (NSNAll pipeline [Robinson et al., 2014]). Finally, Gaussian spatial smoothing was carried out in cortical surface with a Full-Width at Half Maximum of 4mm. In our analysis, 32492 cortical vertices from each subject’s left surface are used.

#### 2.4.2 Evaluation scheme

In the simulation, we consider both linear and non-linear signals. For linear signals, we compare our approach with five other methods: a connectivity-wise general linear model (GLM) approach controlling the family-wise error rate (SPU(Inf) approach [Kim et al., 2014]), principal component regression (PCR) [Jolliffe, 2002], multivariate distance matrix regression (MDMR) [Shehzad et al., 2014], least-squares kernel machine (LSKM) [Liu et al., 2007; Ge et al., 2012] and adaptive sum of powered score (aSPU) and extensions [Kim et al., 2014]. For non-linear signals, we compare our approach with kernel principal component regression (KPCR) [Schölkopf et al., 1997] and LSKM [Liu et al., 2007; Ge et al., 2012]. Among these approaches, only the MDMR approach has previously been used to analyse the dense connectome, but other approaches can achieve a similar goal [Kim et al., 2014].

**Type I error rate:** To evaluate whether an approach can control the type I error rate, we compare two groups of healthy subjects with similar demographic information. It is expected that there is no difference between two groups. Therefore, if a method can provide a valid control, the observed false-positive rate will be around its nominal level 0.05. In detail, first, a voxel is randomly selected, and functional connectivities between it and all other voxels across the whole brain are calculated for every subject. Second, 281 subjects are randomly divided into two groups, and every method is then applied to test whether this voxel is different between the two groups. This step results in a p-value for every approach. Third, the above two steps are repeated for 1000 times, and the observed false positive rate is estimated as the proportion of times the p-value is below 0.05.

**Power:** To compare the power of different methods, a similar method can be adopted but with signals added to real data. This kind of evaluation method makes the situation simulated mimic real data, which has been widely adopted by genome-wide association studies (GWAS) to compare different methods (e.g. [Zhou and Stephens, 2012; Yang et al., 2014]). The power in our simulation is defined as the probability of finding at least one signal (e.g. altered functional connectivity) in a set of functional connectivities, with the false-positive rate being controlled at 0.05.

In our simulation, first, one voxel is randomly selected, and functional connectivities between it and all other voxels across the whole brain are calculated for every subject. Second, signals are then randomly added to a subset of functional connectivities (proportion of null functional connectivity *ρ*). For linear signals, we simply consider the mean difference *γ*_1_ between the two groups. Therefore, the 281 subjects are randomly divided into two groups, and then signals are randomly added to one group. For non-linear signals, we first simulate a Gaussian random noise signal *y*, and then convert it to a four-degree polynomial signal as: *γ*_2_*y*^4^ + 0.5*y*^2^ + 1 and add it to the functional connectivities. Every method is then applied to test the presence of signals. Third, the above two steps are repeated for 1000 times, and the empirical power is estimated by the proportion of times the p-value is below 0.05. In this way, we did not change the underlying connectivity structure (covariance between connectivities).

**Parameter settings and implementation details:** The mean difference parameter *γ*_1_ is set to (0.01,0.02,…, 0.2) and the coefficient of the four-degree polynomial *γ*_2_ is set to (0.001, 0.002,…, 0.05). The proportion of null functional connectivities *ρ* is set to (0.5, 0.75, 0.85, 0.9, 0.95, 0.99). For sKPCR, the top 5 principal components are selected, and the noise covariance matrix is set to the Graph Laplacian operator (based on the neighbourhood relationship between voxels or vertices). For other methods, the MDMR is implemented based on the Matlab script ‘y_CWAS.m’ in Data Processing and Analysis for Brain Imaging (DPABI) [Yan et al., 2016]. Following the paper [Shehzad et al., 2014], the distance matrix for MDMR is calculated as 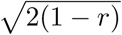, where *r* is the inter-subject correlation matrix. We implement aSPU using the *R* package ‘aSPU’. The power parameters for SPU and aSPU are set to the default values (1, 2,…, 8, Inf). For LSKM, the code was obtained from the author of [Ge et al., 2012].

### 2.5 Real data analysis: brain-wide associations of the schizophrenic connectome

We compared different methods using resting-state fMRI data from the Centers of Biomedical Research Excellence (COBRE) and a Taiwan dataset. In the COBRE dataset, 130 subjects (58 patients with schizophrenia and 72 healthy controls) were used. In the Taiwan dataset, 259 subjects (123 patients with schizophrenia and 136 healthy controls) were used. The detailed data acquisition methods are illustrated in the Appendix B and C. Resting-state fMRI data were preprocessed using the same pipeline as the SWU dataset. Finally, 19567 voxels located in each subject’s brain were extracted for the subsequent analysis.

Our evaluation strategies for sKPCR and other methods include: (1) Evaluating the within-site and between-site reproducibility; (2) Comparing the identified voxel clusters with previously reported schizophrenia findings in the Neurosynth database. For sKPCR only, we also (1) evaluate the robustness of the findings using different model parameters (number of principal components and covariance operators); (2) evaluate the effect of global signal regression; (3) evaluate whether more significant findings really indicate that there exist a larger number of altered functional connections and a better classification accuracy of patients and controls; (4) apply it to identify voxel-clusters that are associated with PANSS score.

For the ease of reading, the details combined with the corresponding results are illustrated in the Results Section together. Throughout the real data analysis, we use the linear sKPCR approach. The proportion of overlap *ρ* used in our analysis is defined as:

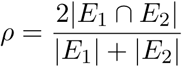

where |*E_i_*| is the number of findings in the *i*-th experiment.

## 3 Results

### 3.1 Simulation study

#### 3.1.1 Type I error rate

We first evaluate whether the all methods can control the type I error rate (see Methods Section 2.4.2 for the detailed evaluation scheme). The results show that all approaches can control the false-positive rate at the nominal level 0.05 in both volume-based and surface-based fMRI data (Figure 2). As our approach is a parametric one, we further evaluate whether the connectivity data, i.e. the Fisher’s Z transformed correlation coefficients, are Gaussian distributed. For each of the 14364 × (14364 − 1)/2 = 103155066 functional connectivities, we use a one-sample Kolmogorov-Smirnov test to test the null hypothesis that the cross-subject normalized data comes from a standard normal distribution. Supplementary Figure D.11 shows that most p-values cannot reject the null hypothesis (p>0.05), which indicate that the normality assumption is met.

**Figure 2:**
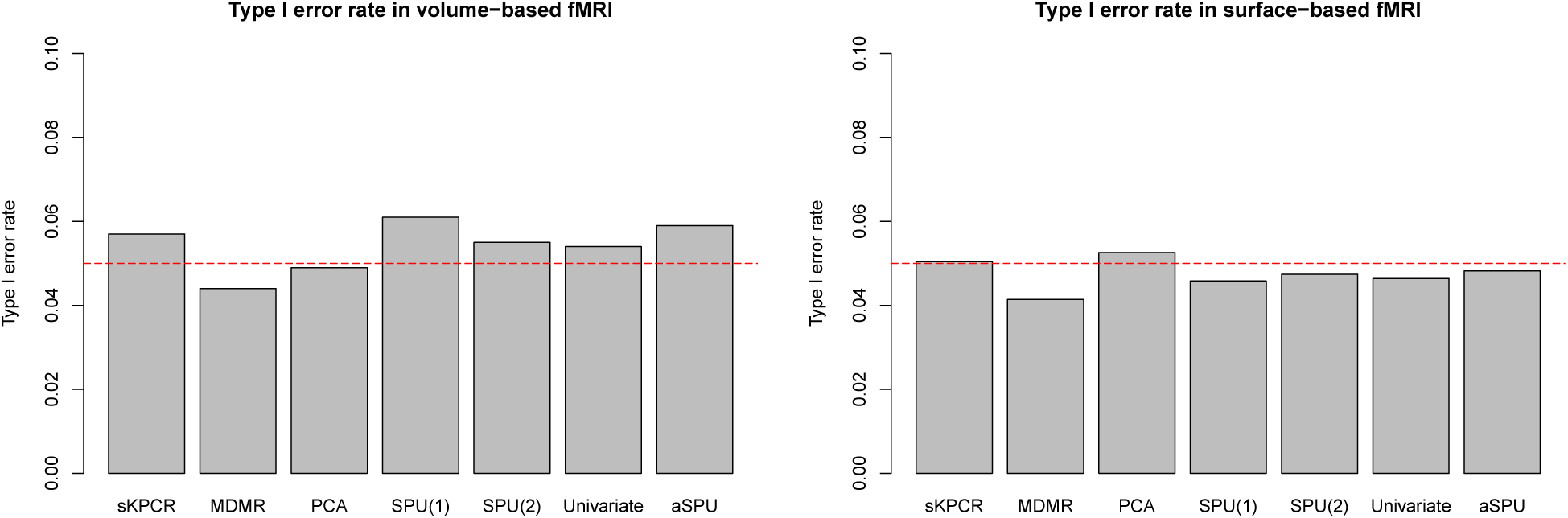
Type I error rate of different methods estimated in volume-based and surface-based fMRI data.

#### 3.1.2 Power comparison

Figure 3 and 4 report the results of comparing the power of different methods when the true signal is *linear,* i.e. the mean difference between the case-control groups (see Methods Section 2.4.2 for the detailed evaluation scheme). For both volume-based and surface-based data, we can clearly observe that the proposed sKPCR method using a linear kernel usually has the highest power in different situations (different effect size and proportion of non-null connectivities). With the proportion of the non-null functional connectivity decreased, the power of multivariate approaches decreased, such as sKPCR, MDMR, PCR, SPU(1) and SPU(2). The power of the univariate approach (GLM) maintains, as it is only related to the connectivity with the maximum effect size. The aSPU approach can be seen as a hybrid approach, thus, it can borrow strength from both multivariate approach and univariate approach.

**Figure 3:**
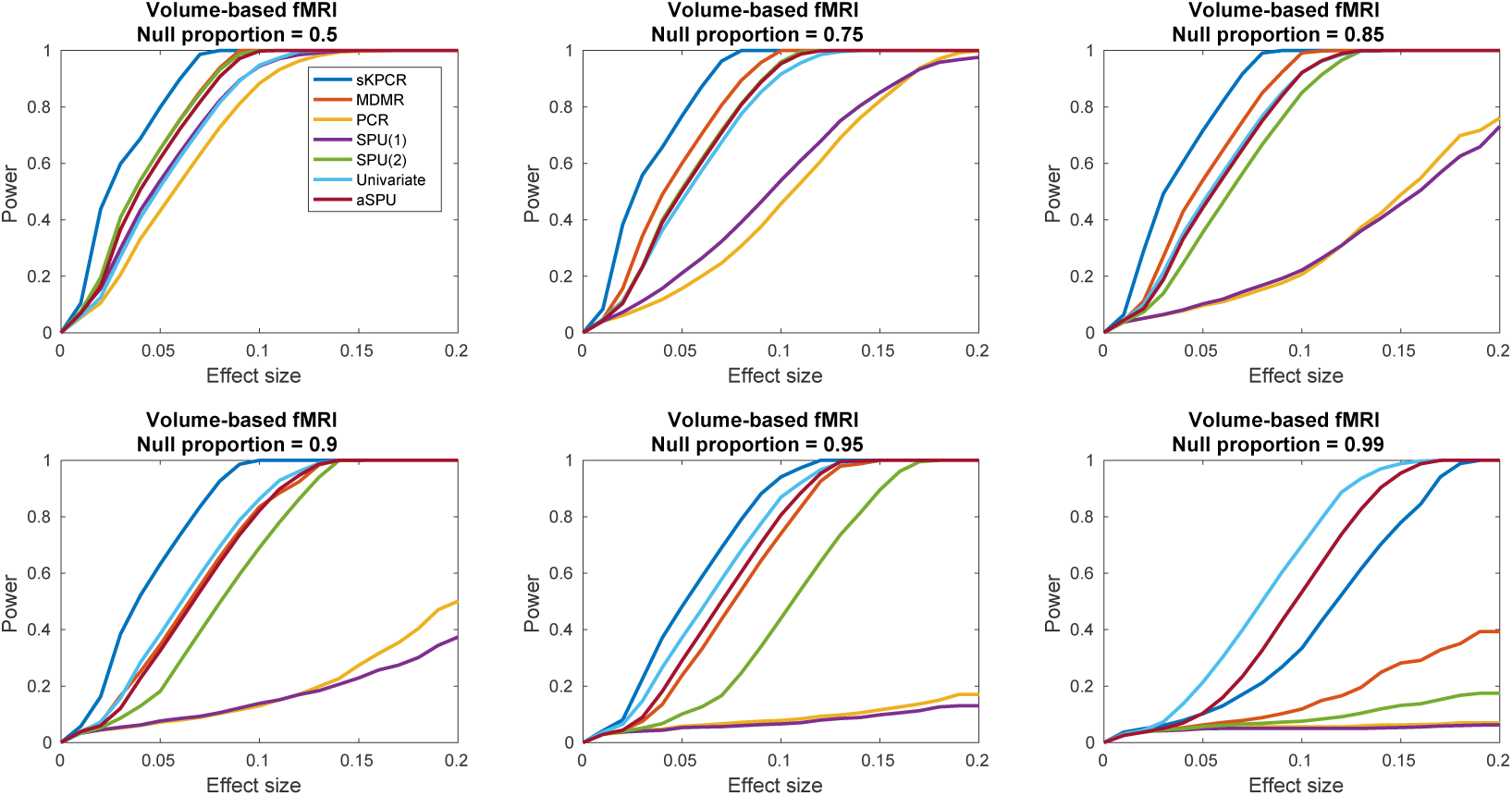
Comparisons of the power of detecting *linear* signals with different methods using simulations relating to *volume-based fMRI data*. Each figure plots the power curves of 7 different methods under different signal effect sizes (between group mean difference = 0.01~0.2) with certain null signal proportions (0.5~0.99).

**Figure 4:**
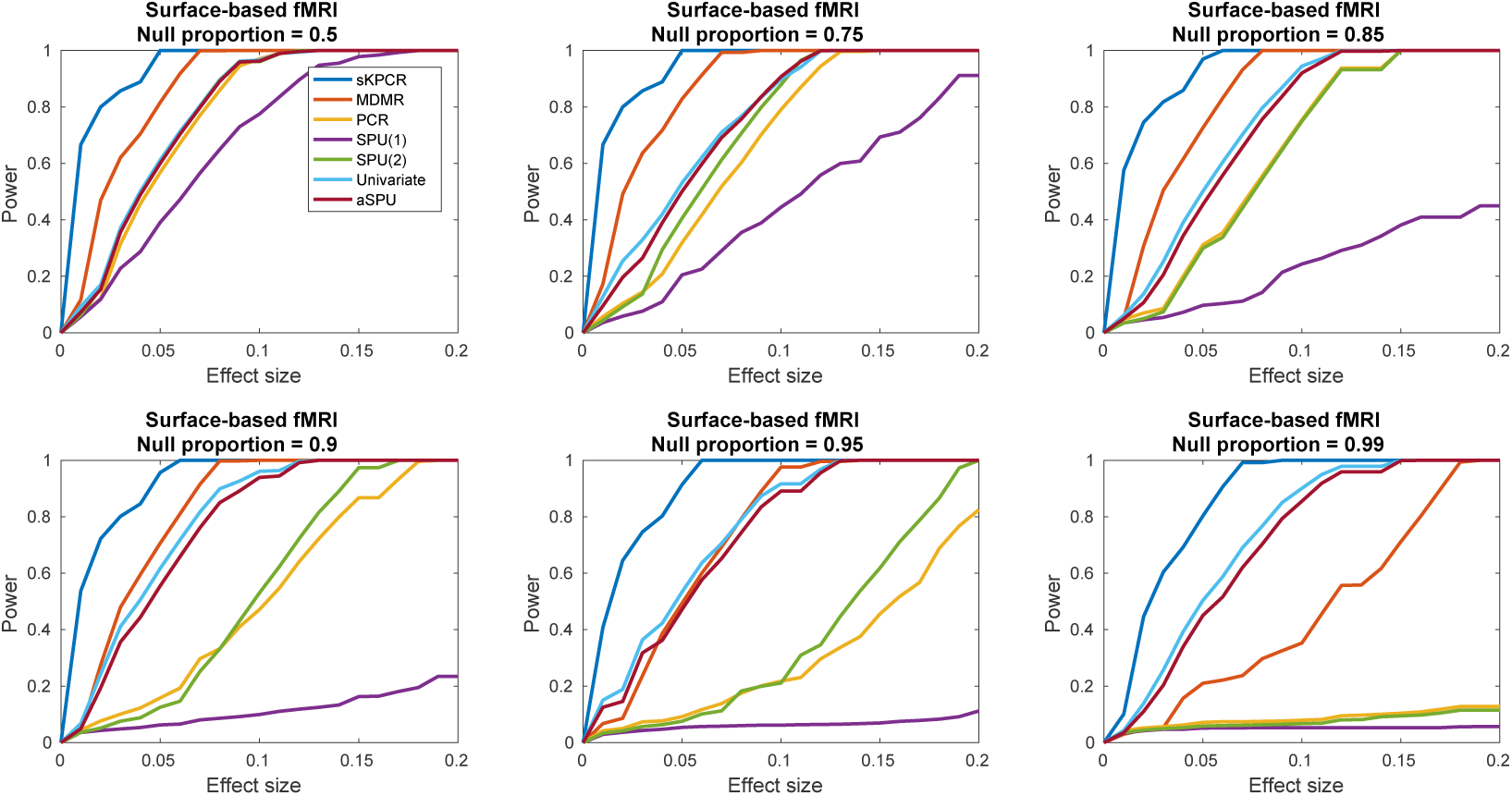
Comparisons of the power of detecting *linear* signals with different methods using simulations relating to *surface-based fMRI data.* Each figure plots the power curves of 7 different methods under different signal effect sizes (between group mean difference = 0.01~0.2) with certain null signal proportions (0.5~0.99).

Figure 5 and 6 report the results of comparing the power of different methods when the true signal is *nonlinear,* i.e. a four-degree polynomial (see Method Section 2.4.2 for the detailed evaluation scheme). Clearly, the sKPCR approach with a 4-degree polynomial kernel always has the highest power. The KPCR usually wins the second place, and the LSKM is usually the third. Compared with the nonlinear methods, other linear methods, such as sPCR, PCR and MDMR, usually have low power.

**Figure 5:**
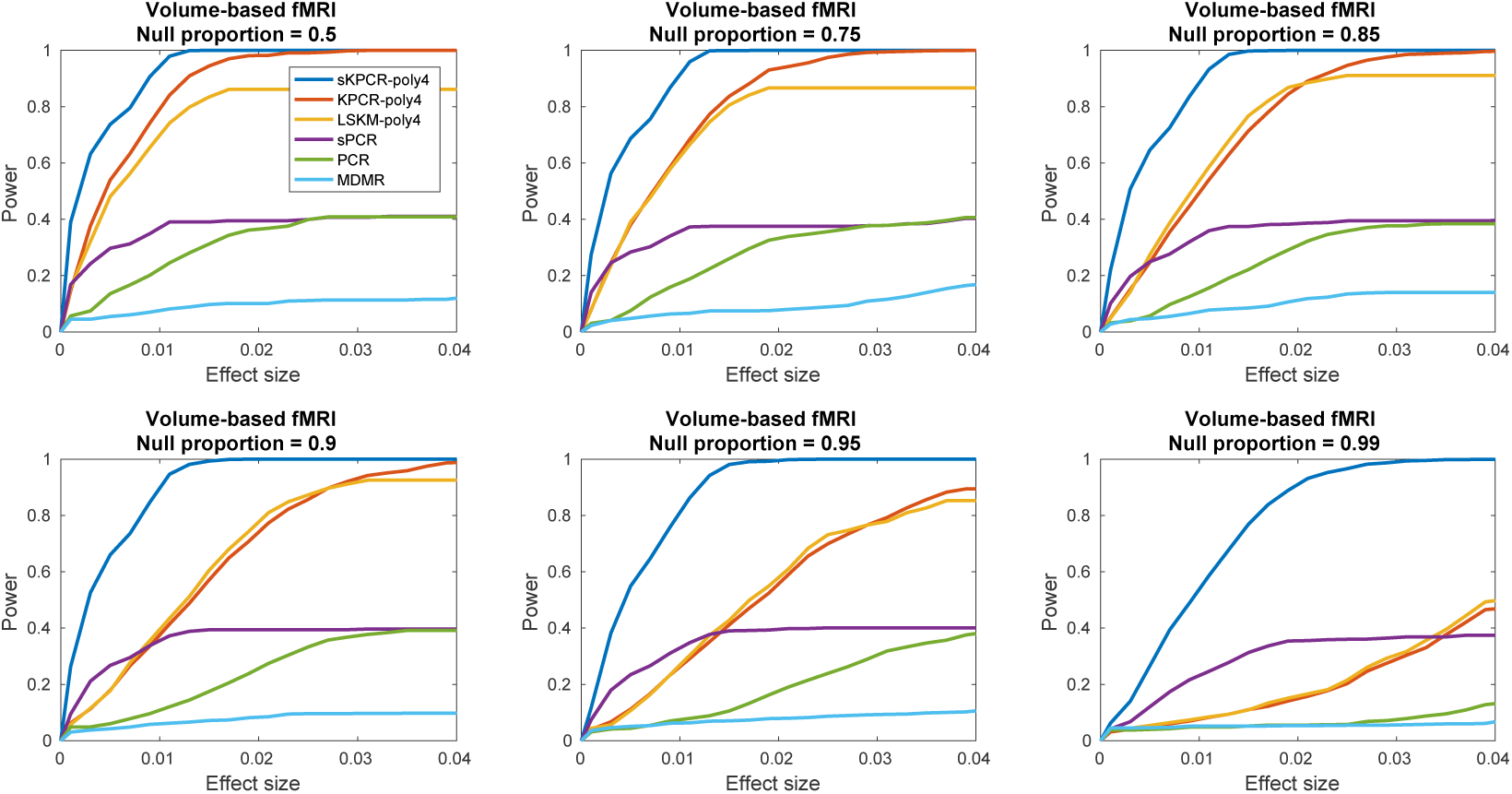
Comparisons of the power of detecting *nonlinear* signals with different methods using simulations relating to *volume-based fMRI data.* The simulated nonlinear signal is a four-degree polynomial γ*y*^4^ + 0.5*y*^2^ +1. Each figure plots the power curves for 6 different methods under different signal effect sizes (γ = 0.001 ~ 0.04) with certain null signal proportions (0.5~0.99).

**Figure 6:**
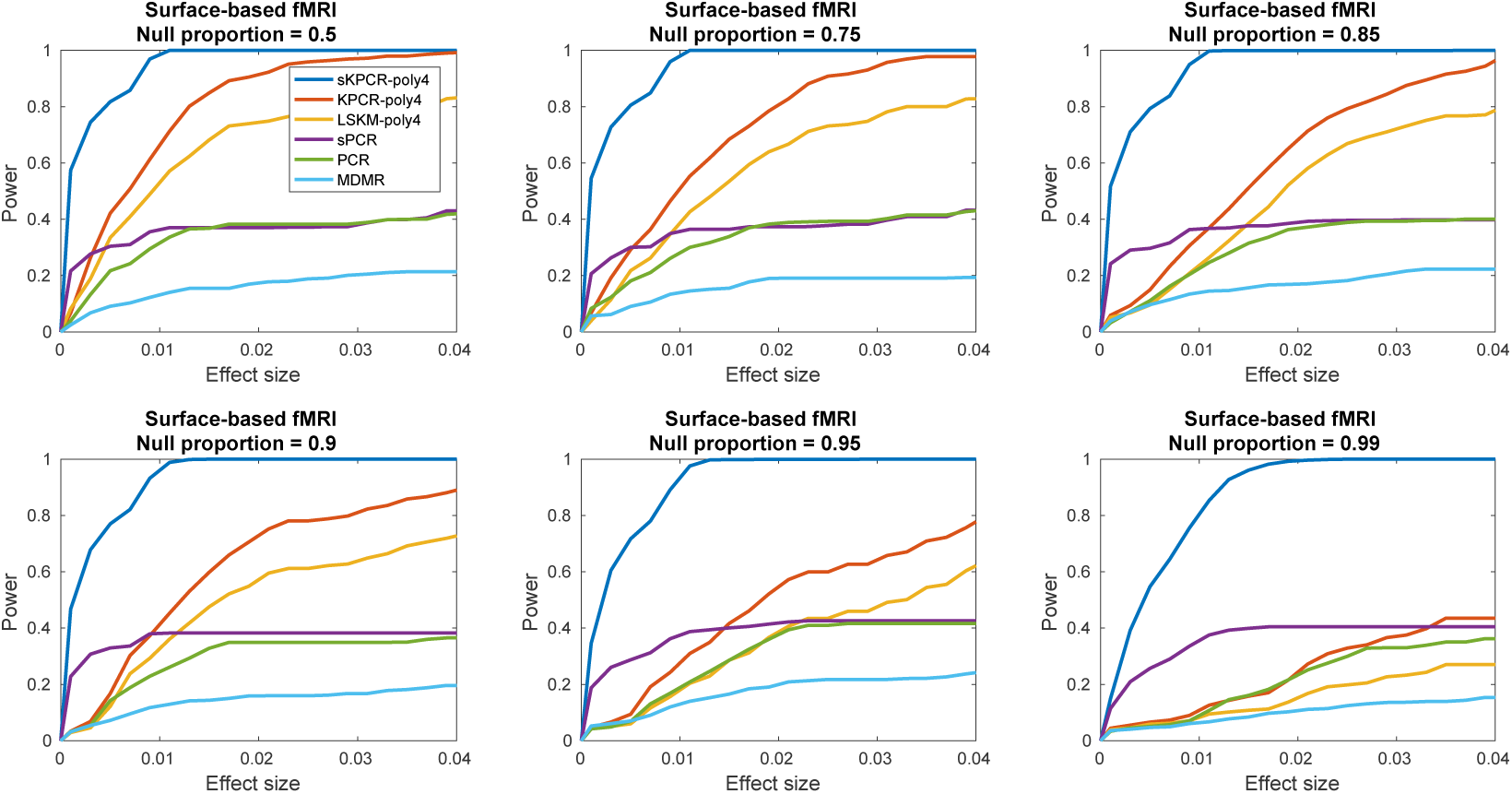
Comparisons of the power of detecting *nonlinear* signals with different methods using simulations relating to *surface-based fMRI data*. The simulated nonlinear signal is a four-degree polynomial γ*y*^4^ + 0.5*y*^2^ +1. Each figure plots the power curves for 6 different methods under different signal effect sizes (γ = 0.001 ~ 0.04) with certain null signal proportions (0.5~0.99).

#### 3.1.3 Computation time

We compare the computation time of sKPCR with MDMR and aSPU using a simple simulation. All the analyses are implemented in Matlab 2016b using a single core on a Linux workstation with Intel Xeon E5-2660 v3(2.60GHz) CPU and 128GB memory. We simulate fMRI data with 27000 voxels/subject. We then compare the computation time for analysing a single voxel using different sample sizes. The time to construct the subject-wise network is omitted, as it is the same across different methods. For MDMR and aSPU, the number of permutations are set to 1000 and 10000. Table 1 shows the results. Our method is the fastest one, and the computation time grows approximately linearly with the sample size. MDMR and aSPU^1^ are much slower, and the computation time grows approximately quadratically with the sample size. When the sample size reaches 500, our method is over 200 times faster than MDMR and over 500 times faster than aSPU using 10000 permutations.

**Table 1:** Comparing the computation time (second) of our method with permutation-based approaches for analysing a single voxel.

| Sample size | 100 | 150 | 200 | 250 | 300 | 350 | 400 | 450 | 500 |
| --- | --- | --- | --- | --- | --- | --- | --- | --- | --- |
| **sKPCR** | **0.27** | **0.28** | **0.32** | **0.34** | **0.38** | **0.40** | **0.44** | **0.51** | **0.55** |
| **MDMR (1000 permutations)** | 0.38 | 0.62 | 1.18 | 1.55 | 2.29 | 3.28 | 3.59 | 5.03 | 7.37 |
| **MDMR (10000 permutations)** | 2.68 | 5.21 | 9.17 | 14.30 | 21.33 | 26.85 | 33.07 | 41.90 | 68.83 |
| **aSPU (1000 permutations)** | 2.96 | 3.39 | 4.03 | 5.97 | 11.09 | 14.24 | 14.69 | 21.59 | 25.74 |
| **aSPU (10000 permutations)** | 30.13 | 34.18 | 45.81 | 60.70 | 95.23 | 114.02 | 142.00 | 149.59 | 168.33 |

**Table 2:** The demographic information of subjects in the COBRE and Taiwan datasets.

| **Dataset** | **Group** | **# Subjects** | **Age (mean ± std)** | **Gender (M/F)** | **mean FD** |
| --- | --- | --- | --- | --- | --- |
| COBRE | Control | 72 | 35.6±11.7 | 50/22 | 0.21±0.11 |
| | Patient | 58 | 36.7±13.5 | 49/9 | 0.24±0.11 |
|  | Statistic (p-value) | NA | 0.61 | 0.07 | 0.07 |
| Taiwan | Control | 136 | 44.1±12.0 | 51/79 | 0.11±0.05 |
| | Patient | 123 | 44.0±11.3 | 51/72 | 0.10±0.07 |
|  | Statistic (p-value) | NA | 0.79 | 1 | 0.66 |

### 3.2 Real data analysis: brain-wide associations of the schizophrenic connectome

#### 3.2.1 Reproducible voxel clusters identified by linear sKPCR

We applied sKPCR to identify voxels whose whole-brain connectivity patterns are different between schizophrenic patients and age and gender matched healthy controls in two resting-state fMRI datasets (COBRE and Taiwan). We used equation (5) to determine the number of principal components at each voxel and graph laplacian operator.

In the COBRE and Taiwan dataset, the proportion of significant voxels is 10.6% (2075 voxels) and 24.0% (4700 voxels) respectively (cluster-based non-parametric permutation test, cluster-defining threshold *p* = 0.001, cluster size *p* < 0.05). Highly similar findings are observed in these two datasets indicating the validity of the proposed method. The proportion of overlap between the two thresholded maps is 31.9% (1082 voxels, p-value of Fisher’s exact test = 4.5 × 10^-323^) and the Pearson correlation coefficient between the two un-thresholded maps is *ρ* = 0.29 (p-value = 0). Figure 7 shows the findings. The five largest overlapped regions between the two datasets are mainly located in the postcentral gyrus, precuneus, thalamus, anterior cingulate cortex and midcingulate cortex ([Tzourio-Mazoyer et al., 2002], Figure 7).

**Figure 7:**
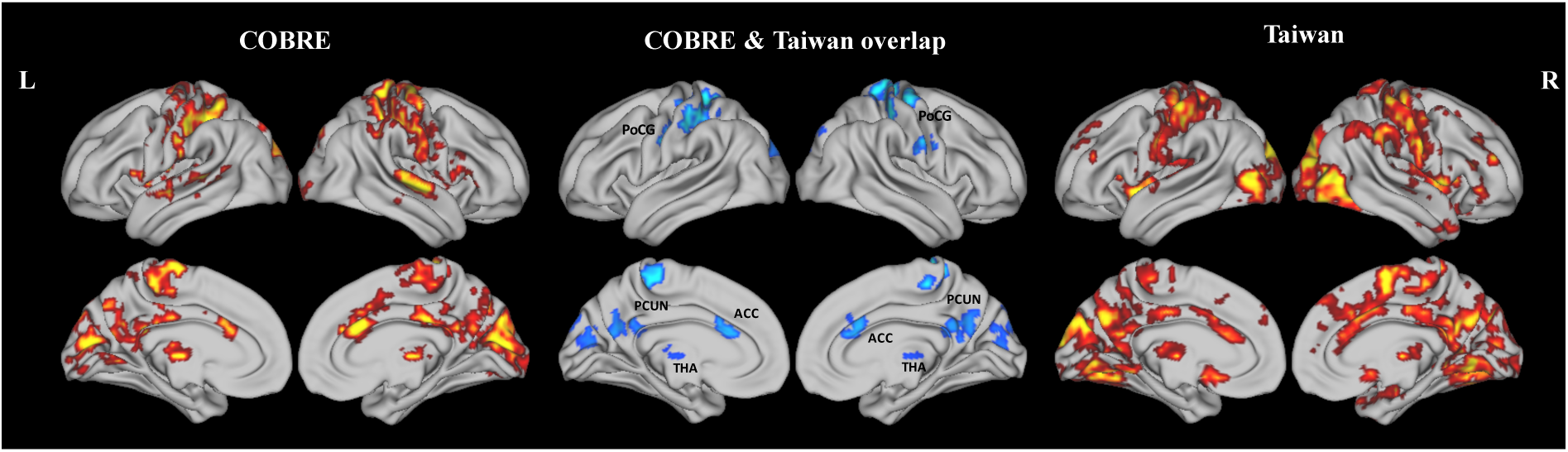
The voxel clusters show significant connectivity pattern differences between schizophrenic patients and healthy controls found by linear sKPCR in the COBRE and Taiwan datasets. The statistical maps are thresholded using permutation-based cluster-wise inference with a cluster-defining threshold of *p* = 0.001 and cluster size *p* < 0.05. Left: the results for the COBRE dataset. Middle: the overlapped clusters for the two datasets. Right: the results for the Taiwan dataset. Abbreviations: PoCG: postcentral gyrus; PCUN: precuneus; THA: thalamus; ACC: anterior cingulate cortex; MCC: midcingulate cortex.

We further performed 1000 bootstrap analyses within each site to test the reproducibility of our method. Figure D.12 shows the results. Again, the results are highly similar between bootstrapped samples within each dataset, with the average proportion of overlap equalling 87% (std = 4%) and 80% (std = 4%) between thresholded maps and average Pearson correlation coefficients equalling 0.46 (std = 0.06) and 0.45 (std = 0.06) in COBRE and Taiwan respectively.

#### 3.2.2 Comparison with other methods

We compared sKPCR with MDMR, SPU(1), SPU(2), univariate (SPU(Inf)) and aSPU methods by applying them to analyse the COBRE and Taiwan datasets. After the statistical maps are obtained, they are all thresholded using the permutation-based cluster-size inference method (cluster-defining threshold *p* = 0.001, cluster size *p* < 0.05). As there is no golden standard, we evaluate them in two ways: (1) Between-sites reproducibility: the proportion of overlap between the findings in the two independent datasets; (2) Literature-based evidence: the proportion of overlap between the findings of different methods and the literature-based meta-analysis findings in the Neurosynth database [Yarkoni et al., 2011]. We used the search term ‘schizophrenia’ for the comparison.

Table 3 shows that our method not only demonstrates a better between-sites reproducibility (32% proportion of overlap between COBRE and Taiwan), but also has a higher proportion of overlap with the existing findings (18% between COBRE and Neurosynth, 29% between Taiwan and Neurosynth). Comparing Figure 7 and Figure 8, it can be clearly observed that the identified regions that overlap with the Neurosynth terms ‘schizophrenia’ are very consistent between the COBRE and Taiwan datasets. They are mainly located in the regions that are reproducible across the two datasets.

**Table 3:** Comparing the reproducibility of different approaches in real data analysis, including between-sites reproducibility and the overlap with the Neurosynth meta-analysis database.

|  | **sKPCR** | **MDMR** | **PCR** | **SPU(1)** | **SPU(2)** | **Univariate** | **aSPU** |
| --- | --- | --- | --- | --- | --- | --- | --- |
| Proportion of overlap (COBRE & Taiwan) | **32%** | 21% | 8% | 2% | 18% | 18% | 22% |
| Proportion of overlap (COBRE & Neurosynth<sup>1</sup>) | **18%** | 7% | 14% | 5% | 6% | 3% | 10% |
| Proportion of overlap (Taiwan & Neurosynth<sup>1</sup>) | **29%** | 15% | 23% | 1% | 13% | 9% | 15% |

<sup>1</sup>The term ‘schizophrenia’ in the Neurosynth database.

**Figure 8:**
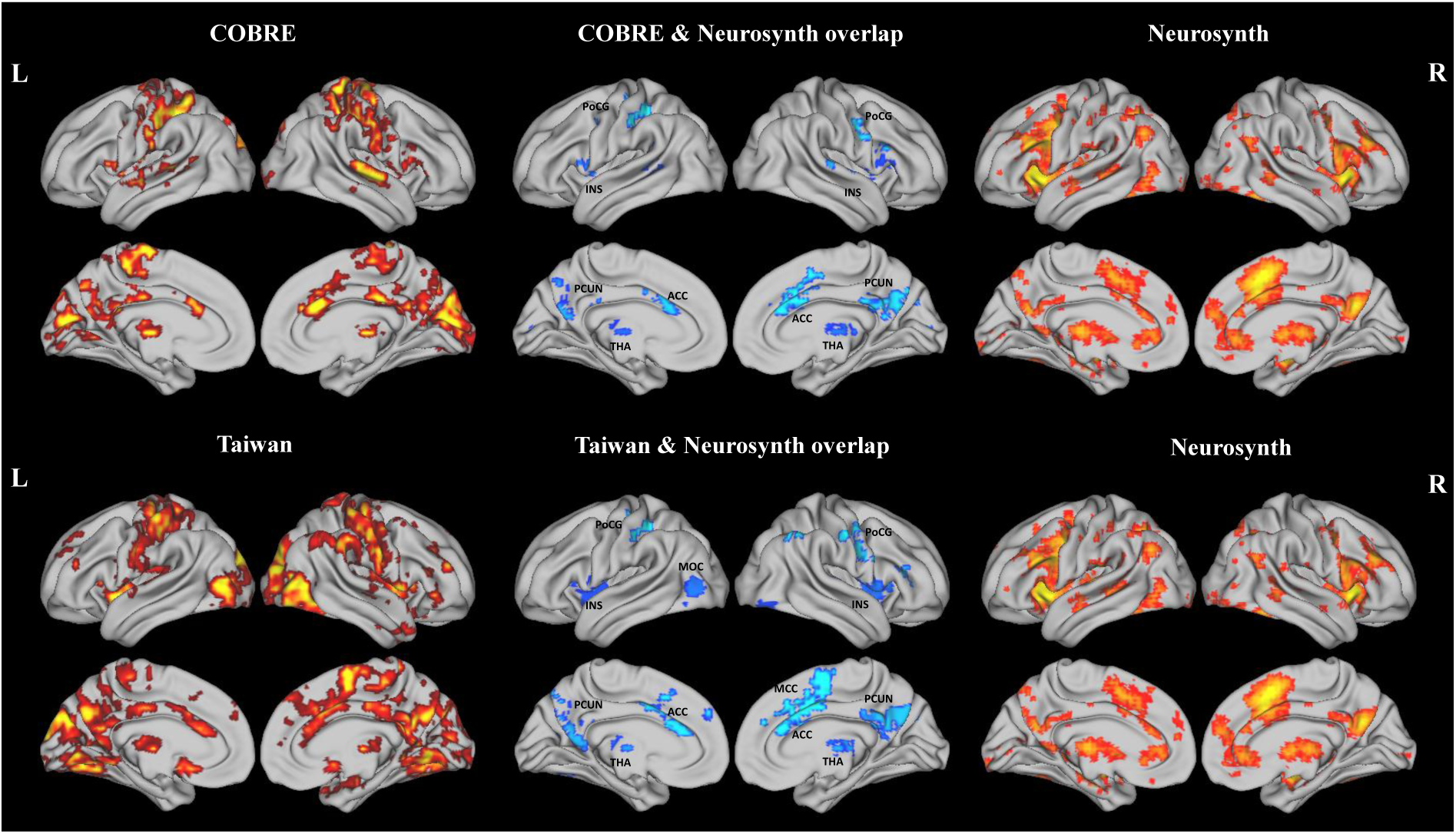
Comparison of the findings of linear sKPCR using the COBRE (upper) and Taiwan (lower) datasets with the activation map produced by the term ‘schizophrenia’ using the Neurosynth database. Left: the results for the two datasets. Middle: the overlapped clusters rom the two datasets. Right (upper and lower): the statistical map for ‘schizophrenia’ using the Neurosynth database. Abbreviations: PoCG: postcentral gyrus; PCUN: precuneus; INS: insular cortex; THA: thalamus; ACC: anterior cingulate cortex; MCC: midcingulate cortex; MOC: middle occipital gyrus.

#### 3.2.3 The stability of sKPCR

In the sKPCR, there are two free parameters that should be pre-specified: the number of principal components and the type of covariance operators. Therefore, to test whether the method is sensitive to the choice of parameters, we performed sKPCR using different parameters in the COBRE dataset. For the number of principal components, we choose the number of PCs from 1 to 50 for comparison. For the covariance operators, we choose the Gaussian Random Field Operator (FWHM = 2,4,6 voxels), Graph Laplacian Operator and Normalized Graph Laplacian Operator for comparison. The sKPCR was applied to analyse the COBRE dataset with different combinations of the two parameters, and the correlation coefficients between the pairwise raw statistical maps (− log_10_(p-value)) were calculated. Figure 9 shows that the results obtained using different parameters are highly similar to each other. Most of the time, the correlation coefficients are larger than 0.95. The results indicate that sKPCR is insensitive to model parameters. Actually, we find that the significant regions found using different parameters are highly similar in the real data analyses (the results are not shown here due to limited space).

**Figure 9:**
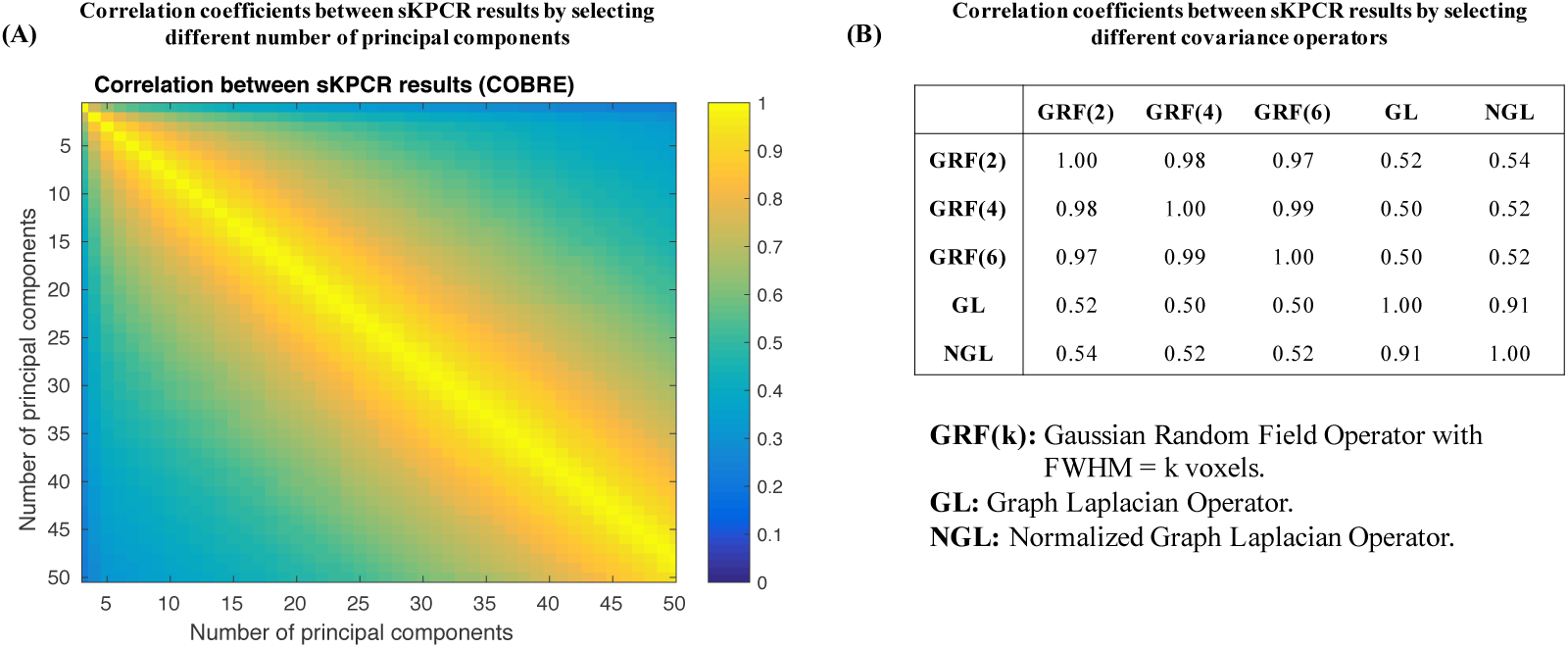
The Pearson correlation coefficients between the sKPCR results using different model parameters in the COBRE dataset. The correlation was computed between two raw -log_10_(p) statistical maps. (A) Selecting different number of principal components (from 1 to 50) at each voxel; (B) Selecting different covariance operators, including: the Gaussian Random Field Operator (FWHM = 2, 4, 6 voxels), Graph Laplacian Operator, and the Normalized Graph Laplacian Operator.

The preprocessing pipeline may also affect the results of sKPCR, for example, global signal regression (GSR). At present, there is no consensus whether or not to regress out the global signal before computing the subject-level connectivity matrix. Therefore, we examined the effect of GSR in our analysis. Despite the percentage of significant findings being higher without GSR (6170 and 7656 significant voxels in the COBRE and Taiwan datasets at CDT=0.001 and cluster size *p* < 0.05), the results are highly correlated with the results with GSR within each site (Taiwan *ρ* = 0.73, COBRE *ρ* = 0.67). The key clusters identified by sKPCR are highly similar, including, for example, the thalamus, Postcentral gyrus, Insula, Precuneus, Cuneus and Mid Cingulate cortex (Supplementary Figure D.13). In sum, the sKPCR is highly stable across different parameter settings and preprocessing pipelines.

#### 3.2.4 Connection with the connectivity-wise univariate approach

We also examined whether the voxels with a more significant p-value connect a larger number of different functional connectivities. We applied the connectivity-wise general linear model (GLM) analysis to both datasets. The same nuisance covariates, including age, sex, mean FD, are used in the GLM analysis. The GLM result is a Z-statistic at each connectivity. To summarize the connectivity-wise results at the voxel level, we calculate, for each voxel, the sum of Z^2^ of all the connectivities connecting this voxel across the whole brain. We can observed a strong positive correlation between the sum of Z^2^ and the −log_10_(p-value) of sKPCR per voxel (Figure 10). For the COBRE dataset, the correlation coefficient is 0.671, and for the Taiwan dataset, it is 0.698.

**Figure 10:**
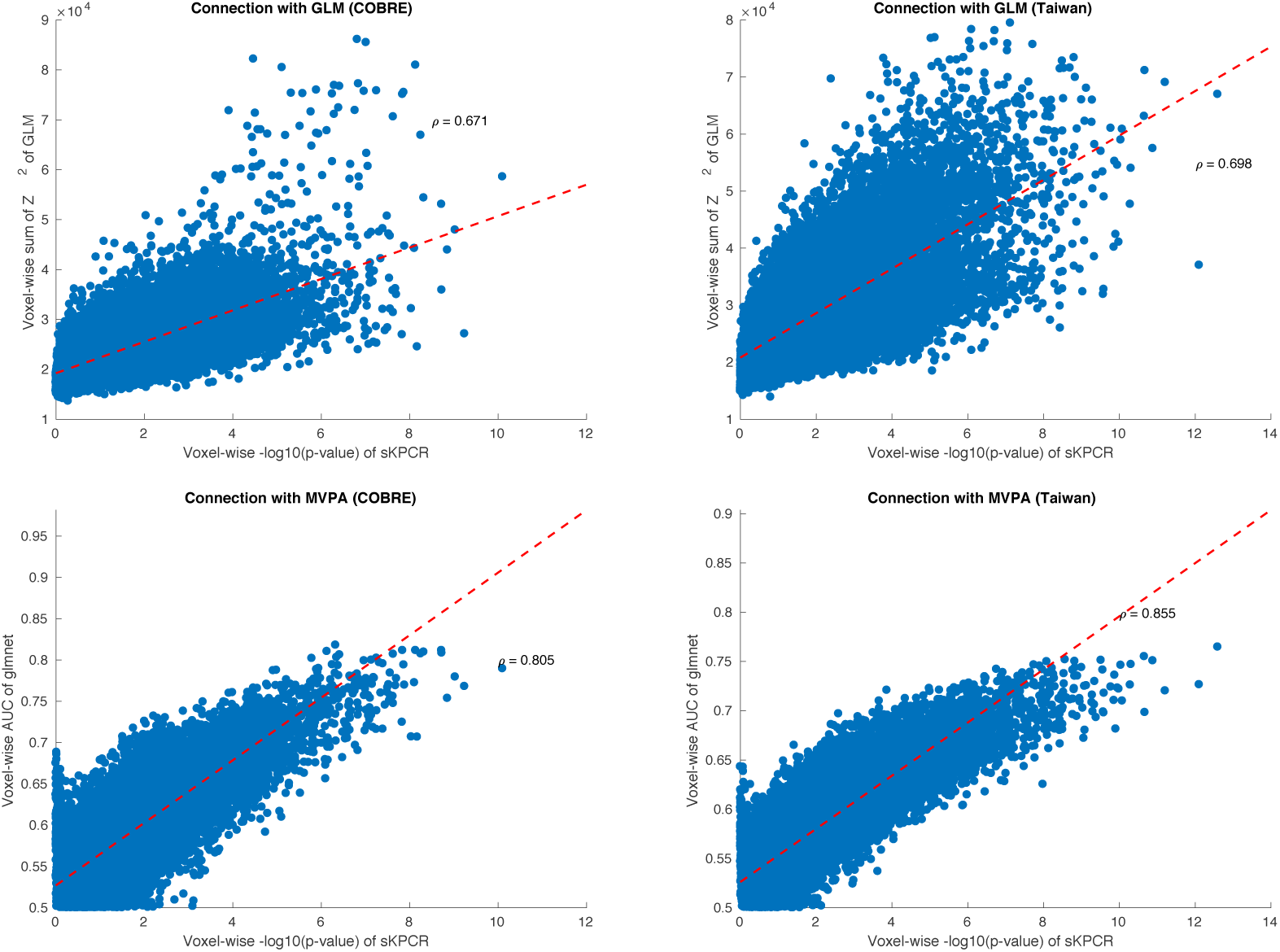
Comparing sKPCR with other methods in the COBRE and Taiwan dataset. **Top row:** Comparing sKPCR with connectivity-wise GLM analysis. For each figure, a voxel’s significance (− log_10_(p-value)) based on the sKPCR analysis (x-axis) is plotted against the sum of squares of the Z-statistics of individual connections for the same voxel in a GLM analysis (y-axis). **Bottom row:** Comparing sKPCR with MVPA. A voxel’s significance (− log_10_(p-value)) based on the sKPCR analysis (x-axis) is plotted against the ten-fold cross validation AUC using a elastic-net regression to classify patients and controls for the same voxel (y-axis).

#### 3.2.5 Connection with voxel-wise multivariate pattern analysis

Finally, we examined whether the voxels with a more significant p-value have a better ability to distinguish schizophrenic patients and healthy controls in a classification task. For each voxel, we applied the widely-used elastic-net regression approach [Zou and Hastie, 2005; Friedman et al., 2010] to classify the subjects. The number of principal components used was the same as the standard sKPCR analysis (equation (5)). A 10-fold cross-validation was performed and the area under the curve (AUC) was used to measure the degree of discrimination between the two populations at each voxel. We observed a strong positive correlation between the AUC and the −log_10_(p-value) of sKPCR per voxel (Figure 10). For the COBRE dataset, the correlation coefficient is 0.805, and for the Taiwan dataset, it is 0.855.

### 3.3 Application of sKPCR to continuous variable phenotypes

We further applied sKPCR to identify clusters that are associated with the Positive and Negative Syndrome Scale (PANSS) score. A total of 181 schizophrenic patients in these two datasets were pooled together (58 in COBRE and 123 in Taiwan), and sKPCR was applied to associate each voxel with the PANSS positive and negative scale. Each score was positive by summing the corresponding sub-item scores. As the PANSS scores were discrete variables which may not be normally distributed, we applied a Box-Cox transformation to improve the normality of the data. The age, gender, sites, and motion (mean FD) were treated as nuisance covariates in our analysis. We used a permutation-based cluster size inference method to identify significant clusters (with cluster-defining threshold *p* = 0.01, cluster size *p* = 0.05). The results are shown in Figure D.14, Figure D.15 and Table 4.

**Table 4:** Brain-wide associations identified by sKPCR in the COBRE and Taiwan datasets.

| Overlapped clusters between the COBRE and Taiwan datasets |  |  |  |  |  |  |  |  |
| --- | --- | --- | --- | --- | --- | --- | --- | --- |
| Clusters | Areas in AAL template | # voxels | Peak statistic<br>(-log10(p)) | MNI coordinates |  |  | # decreased FC<br>in SCZ <sup>1</sup> | # increased FC<br>in SCZ <sup>1</sup> |
| 1 | Cuneus_L, Cuneus_R, Occipital_Sup_L | 131 | 7.29 | -8 | -76 | 20 | 1316 | 746 |
| 2 | Postcentral_R, Precentral_R | 125 | 6.64 | -48 | -24 | 44 | 3811 | 2536 |
| 3 | Postcentral_L, Precentral_L | 124 | 6.73 | 36 | -16 | 64 | 4256 | 1502 |
| 4 | Thalamus_L, Thalamus_R | 98 | 7.87 | 8 | -16 | 4 | 9362 | 21505 |
| 5 | Precuneus_L, Cingulate_Post_L | 75 | 6.36 | -4 | -56 | 24 | 131 | 126 |
| 6 | Insula_L, Putamen_L | 73 | 6.41 | 36 | 12 | 8 | 5042 | 584 |
| 7 | Cingulate_Ant_L, Cingulate_Mid_L | 42 | 5.68 | 4 | 20 | 28 | 730 | 167 |
| 8 | Postcentral_L | 17 | 6.86 | 60 | -8 | 24 | 1402 | 387 |
| Clusters associated with the PANSS positive symptoms |  |  |  |  |  |  |  |  |
| Clusters | Areas in AAL template | # voxels | Peak statistic<br>(-log10(p)) | MNI coordinates |  |  | # positively<br>correlated FC <sup>2</sup> | # negatively<br>correlated FC <sup>2</sup> |
| 1 | Frontal_Inf_Tri_L, Frontal_Mid_L, Frontal_Inf_Oper_L | 152 | 5.10 | 44 | 28 | 44 | 1136 | 978 |
| 2 | Temporal_Inf_L, Temporal_Mid_L, Temporal_Sup_L | 121 | 5.03 | 44 | -4 | -12 | 534 | 218 |
| 3 | Cingulate_Mid_L, Supp_Motor_Area_L | 90 | 4.24 | 4 | -8 | 36 | 322 | 395 |
| 4 | Frontal_Mid_R, Frontal_Inf_Tri_R | 77 | 6.10 | -40 | 24 | 36 | 282 | 266 |
| 5 | Precuneus_L, Parietal_Sup_L | 57 | 3.85 | 28 | -44 | 72 | 63 | 118 |
| 6 | Occipital_Mid_L, Occipital_Inf_L | 50 | 4.93 | 32 | -88 | 0 | 556 | 220 |
| 7 | Cingulate_Ant_L, Cingulate_Mid_L | 42 | 4.44 | 4 | 28 | 28 | 156 | 79 |
| 8 | Temporal_Pole_Sup_R, Temporal_Pole_Mid_R | 41 | 4.59 | -36 | 20 | -32 | 193 | 324 |
| 9 | Angular_R, Parietal_Inf_R | 38 | 4.00 | -48 | -52 | 36 | 94 | 86 |
| 10 | Occipital_Inf_R | 23 | 5.51 | -44 | -68 | -20 | 122 | 88 |
| Clusters associated with the PANSS negative symptoms |  |  |  |  |  |  |  |  |
| Clusters | Areas in AAL template | # voxels | Peak statistic<br>(-log10(p)) | MNI coordinates |  |  | # positively<br>correlated FC <sup>2</sup> | # negatively<br>correlated FC <sup>2</sup> |
| 1 | Temporal_Mid_L, Temporal_Sup_L, Frontal_Inf_Oper_L | 475 | 5.95 | 40 | 12 | 28 | 2154 | 3077 |
| 2 | Pallidum_R, Putamen_R, Insula_R | 133 | 5.45 | -28 | -8 | 4 | 600 | 453 |
| 3 | Frontal_Mid_L, Frontal_Sup_L, Frontal_Inf_Tri_L | 106 | 4.17 | 36 | 60 | 16 | 376 | 530 |
| 4 | Postcentral_L, Precentral_L | 69 | 4.34 | 48 | -28 | 60 | 199 | 373 |
| 5 | Precuneus_R, Occipital_Sup_R | 62 | 4.43 | -20 | -64 | 48 | 510 | 196 |
| 6 | Precentral_R, Postcentral_R | 53 | 3.90 | -56 | -8 | 40 | 480 | 507 |
| 7 | Frontal_Sup_Medial_L, Frontal_Sup_L | 40 | 3.80 | -4 | 40 | 44 | 332 | 153 |
| 8 | Postcentral_R, Precuneus_R | 37 | 4.17 | -28 | -40 | 56 | 141 | 137 |
| 9 | Lingual_L, Occipital_Inf_L | 33 | 3.55 | 16 | -84 | -4 | 87 | 56 |
| 10 | Angular_L | 27 | 3.64 | 40 | -72 | 32 | 73 | 53 |

<sup>1</sup>FCs with p-values smaller than $1 \times 10^{-6}$ .<sup>2</sup>FCs with p-values smaller than $1 \times 10^{-4}$ .

## 4 Discussion

### 4.1 Summary of the proposed method

The sKPCR proposed in this paper is a powerful and efficient approach for dense connectome analysis. Its utilities are demonstrated in both volume-based and surface-based fMRI data for both linear and non-linear signals. It adopts a newly developed sKPCA approach for voxel-wise feature extraction in the connectome. The sKPCA can extract either linear or non-linear signals in the connectome efficiently, while accounting for the specific structure of the dense connectome. Therefore, such a suitable dimension reduction approach makes subsequent statistical analysis more powerful.

A voxel/vertex identified by this approach can be interpreted as ‘there may exist one or more functional connectivities which connect it that are associated with the phenotype of interest’. If one wishes to know the associated connections, a subsequent seed-based analysis can be performed. That is, we can extract a seed time series by averaging the voxel/vertex’s time series within a significant cluster, and test the associations between the seed connectivity map and a phenotype of interest. However, one may not find any significant individual connections in the seed-based analysis, because our approach can detect more than this kind of signal. For example, consider a scenario that many of the connections only have small effect sizes. In addition, as our approach can produce a voxel-wise statistical map, but not a connectivity-wise result, it can be directly compared with results of other analyses, such as task-activation studies, VBM analysis, and Neurosynth meta-analysis results, though our result does not have a direction.

The BWAS approach is another approach for dense connectome association study. It combines connectivity-wise univariate tests and random field theory [Gong et al., 2017]. Conventional topological inference methods, such as peak- and cluster-level inference, are generalized to analyse the dense connectome. It can test the odds that either effect size of every single functional connectivity (peak-level inference) or the spatial extent of functional connectivity clusters exceeding a cluster-defining threshold (CDT) (cluster-level inference) is large by chance. Although we have demonstrated that the sKPCR is more powerful than the peak-level inference approach, it is very hard to compare the power of sKPCR and the cluster-level inference approach proposed by [Gong et al., 2017]. This is because they have different purposes: the sKPCR is to identify voxels, while the cluster-level inference is to identify functional connectivity clusters.

With an increasing popularization of network-based analysis in the neuroimaging field, we believe that a powerful approach is required for the identification of the associations, especially for ultra highdimensional dense connectome data. With more and more high quality fMRI data available, such as the Human Connectome Project [Glasser et al., 2016] and UK Biobank [Miller et al., 2016], it can be expected that there will be a trend from the region-of-interest studies to dense connectome association studies, because the dense connectome can provide more precise locations of the variations in the brain. Recent studies have demonstrated that such dense connectome analyses can precisely identify altered connectivity patterns in several mental disorders, such as schizophrenia [Cheng et al., 2015a], autism [Cheng et al., 2015b] and depression [Cheng et al., 2016; Satterthwaite et al., 2016]. Notably, for major depressive disorder, the identified ‘non-reward’ circuit centered on the right lateral orbitofrontal cortex (OFC) by [Cheng et al., 2016] has shown efficacy in rTMS treatment of depression [Fettes et al., 2017].

### 4.2 Biological significance of the schizophrenia findings

In the present study, voxels with different functional connectivity was found in schizophrenia involving a number of brain areas as shown in Figure D.14, Figure D.15 and Table 4. The regions included the precuneus and cuneus, the anterior, mid, and posterior cingulate cortex; the pre- and post-central gyrus; and the thalamus (Table 4). It was also possible to reveal correlations with the positive symptoms of schizophrenia functional connectivity involving the inferior frontal gyrus; the temporal cortex; the cingulate cortex, the angular gyrus; and the precuneus (Table 4 and Figure D.14). It was also possible to reveal correlations with the negative symptoms of schizophrenia of functional connectivity involving the inferior frontal gyrus; the temporal cortex; the cingulate cortex, the angular gyrus; the precuneus, the frontal cortex; and the pre- and postcentral gyrus (Table 4 and Figure D.15). These correlations with the negative symptoms help to reveal the power of the present approach, for correlations of functional connectivity with the negative symptoms do not always emerge, even though the negative symptoms provide the main source of variation between patients with schizophrenia [Rolls et al., 2017].

These regions have in general been implicated in schizophrenia previously, as shown by the comparison with the results from the Neurosynth database illustrated in Figure 8. Previous research on resting state functional connectivity in schizophrenia has shown widespread functional disconnectivity in distributed brain networks in schizophrenia [Khamsi, 2012; Northoff and Duncan, 2016]. However, very consistent patterns and principles of altered connectivity in schizophrenia remain somewhat elusive [Meyer-Lindenberg, 2010; Whitfield-Gabrieli and Ford, 2012; Northoff and Duncan, 2016]. In one study, first-episode patients had many differences in functional connectivity involving the inferior frontal gyri (Broca’s area), and these changes were correlated with delusions/blunted affect [Li et al., 2017a]. For chronic patients, functional-connectivity differences extended to wider areas of the brain, including reduced thalamo-frontal connectivity, and increased thalamo-temporal and thalamo-sensorimoter connectivity that were correlated with the positive, negative, and general symptoms, respectively [Li et al., 2017a].

The present approach is powerful in detecting voxels with different functional connectivity in schizophrenia. To analyse which functional connectivities are increased or decreased in schizophrenia, the next step having identified the voxels is to use these voxels in a seed-based functional connectivity approach, as described elsewhere in this paper. We leave that seed-based approach for future research, as the main aim of the current research is to describe this new approach.

### 4.3 Limitations and further areas for refinements

There also exist further areas for refinement. First, the method can be extended to analyse multimodal data. For example, a joint analysis of structure and functional networks at either the voxel or ROI level. Extending the ideas of sKPCR to some popular data fusion methods, such as canonical correlation analysis (CCA) and its extensions, should be a possible choice. A comprehensive study should be conducted in the future to validate such methods. Second, the current sKPCR method is designed for single-site studies. Therefore, combining the sKPCR results from multiple imaging sites is an important issue in the future. Possible methods include conventional meta-analysis methods and the model-based site-effect adjustment methods, such as ComBat [Johnson et al., 2007]. Fourth, future investigations can apply the sKPCR approach to more phenotype variables and more publicly available datasets, such as HCP and the UK Biobank, to provide a more comprehensive understanding of the approach, including looking for the best model parameters and the best voxel-wise multiple correction approaches.

## 7 Appendix

### A Data Acquisition: Southwest University

In resting-state fMRI scanning, the subjects were instructed to rest without thinking about a particular topic, and not to fall asleep or close their eyes. The 8-min scan of 281 contiguous whole-brain resting-state functional images was obtained using gradient-echo planar imaging (EPI) sequences with the following parameters: slices = 32, repetition time (TR)/echo time (TE) = 2000/30 ms, flip angle = 90, field of view (FOV) = 220 mm × 220 mm, and thickness/slice gap = 3/1 mm, voxel size 3.4 × 3.4 × 3 mm^3^. A magnetization-prepared rapid gradient echo (MPRAGE) sequence was used to acquire high-resolution T1-weighted anatomical images (repetition time = 1900 ms, echo time =2.52 ms, inversion time = 900 ms, flip angle = 90 degrees, resolution matrix = 256 × 256, slices = 176, thickness =1.0 mm, voxel size = 1 × 1 × 1 mm^3^.

### B Data Acquisition: COBRE

COBRE contributed raw anatomical and functional MR data from 72 patients with Schizophrenia and 75 healthy controls (ages ranging from 18 to 65 in each group). All subjects were screened and excluded if they had: a history of neurological disorder, history of mental retardation, history of severe head trauma with more than 5 minutes loss of consciousness, history of substance abuse or dependence within the last 12 months. Diagnostic information was collected using the Structured Clinical Interview used for DSM Disorders (SCID).

A multi-echo MPRAGE (MEMPR) sequence was used with the following parameters: TR/TE/TI = 2,530/[1.64, 3.5, 5.36, 7.22, 9.08]/900 ms, flip angle = 7°, FOV = 256 × 256 mm, Slab thickness = 176 mm, Matrix = 256 × 256 × 176, Voxel size =1 × 1 × 1 mm^3^, Number of echos = 5, Pixel bandwidth =650 Hz, Total scan time = 6 min. With 5 echoes, the TR, TI and time to encode partitions for the MEMPR are similar to that of a conventional MPRAGE, resulting in similar GM/WM/CSF contrast. Resting-state fMRI data was collected with single-shot full *k*-space echo-planar imaging (EPI) with ramp sampling correction using the intercomissural line (AC-PC) as a reference (TR: 2 s, TE: 29 ms, matrix size: 64 × 64, 32 slices, voxel size: 3 × 3 × 4 mm^3^).

### C Data Acquisition: Taiwan

MRI data were acquired using a 3T MR system (Siemens Magnetom Tim Trio) at National Yang-Ming University, Taipei, Taiwan, equipped with a high-resolution 12-channel head array coil. To minimize head motion, each subject’s head was immobilized with cushions inside the coil during scanning. An anatomical T1-weighted image was acquired with a sagittal 3D magnetization-prepared rapid gradient echo sequence: repetition time (TR) = 3,500 ms, echo time (TE) = 3.5 ms, voxel size = 1 × 1 × 1 mm^3^. Resting-state fMRI data were acquired while subjects were lying quietly and with their eyes closed in the scanner, using a gradient echo-planar imaging sequence sensitive to blood oxygenation level dependent contrast: TR = 2,500 ms, TE = 27 ms, flip angle = 90°, FOV = 220 × 220 mm^2^, thickness = 3.4 mm without gap, matrix size = 64 × 64, voxel size = 3.44 × 3.44 × 3.4 mm^3^, 200 volumes.

### D Supplementary Tables and Figures

**Figure D.11:**
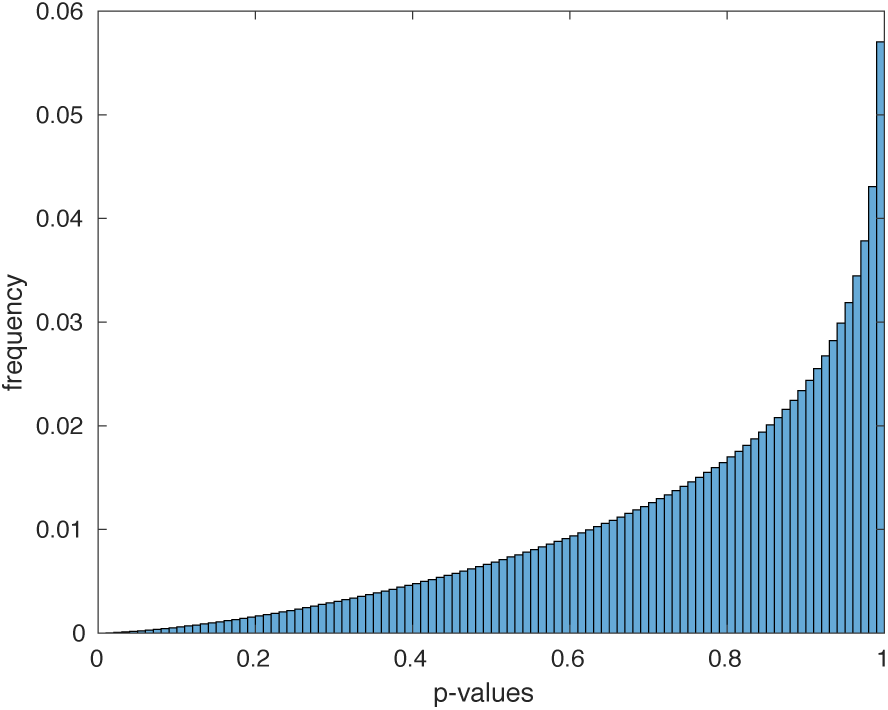
A p-value histogram for the test of normality. For almost all the functional connectivities the null hypothesis that they are Gaussian distributed (p>0.05) can not be rejected.

**Figure D.12:**
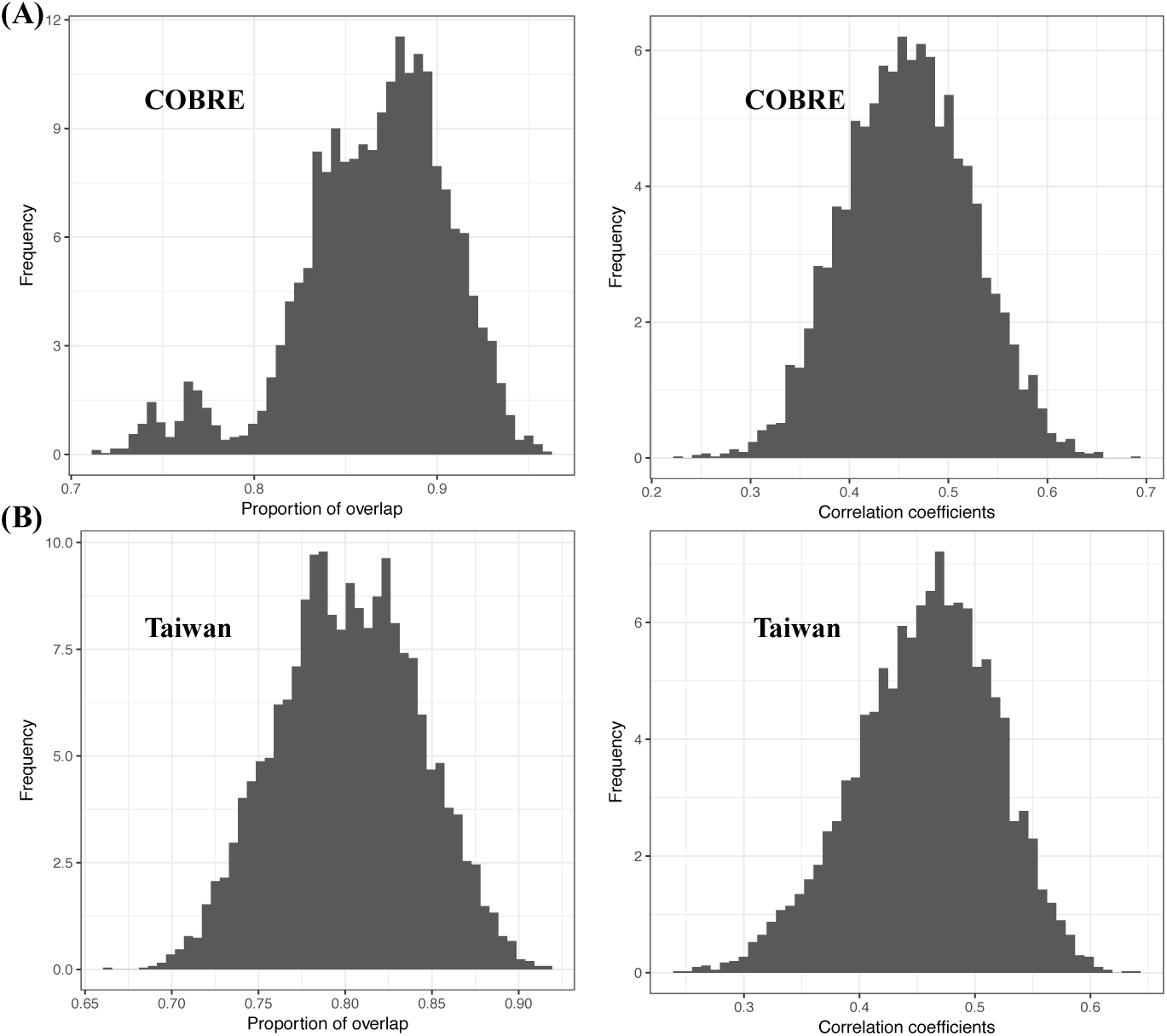
The bootstrap distribution of the proportion of overlap between the thresholded map and the correlation coefficients between the unthresholded map in (A) the COBRE and (B) the Taiwan datasets.

**Figure D.13:**
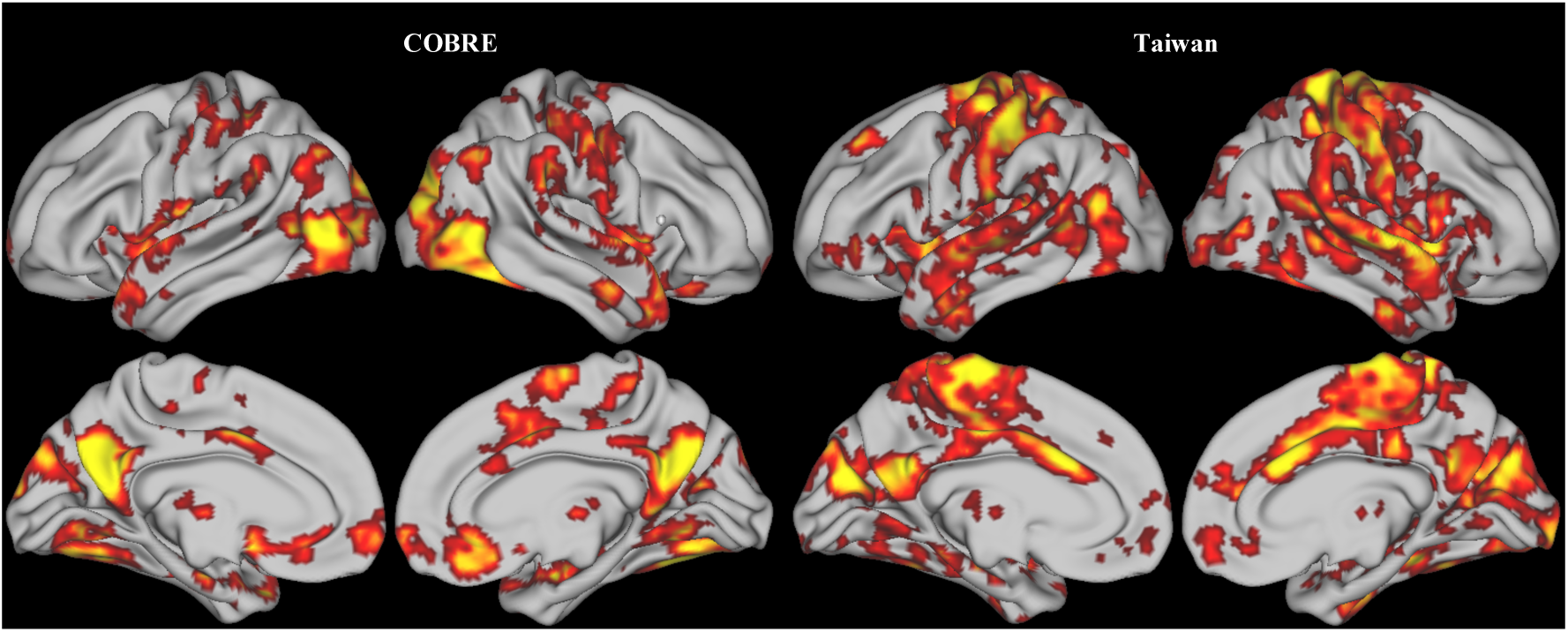
The voxel clusters showing significant connectivity pattern difference between schizophrenic patients and healthy controls found by the linear sKPCR in the COBRE dataset (left) and Taiwan dataset (right) **without global signal regression**.

**Figure D.14:**
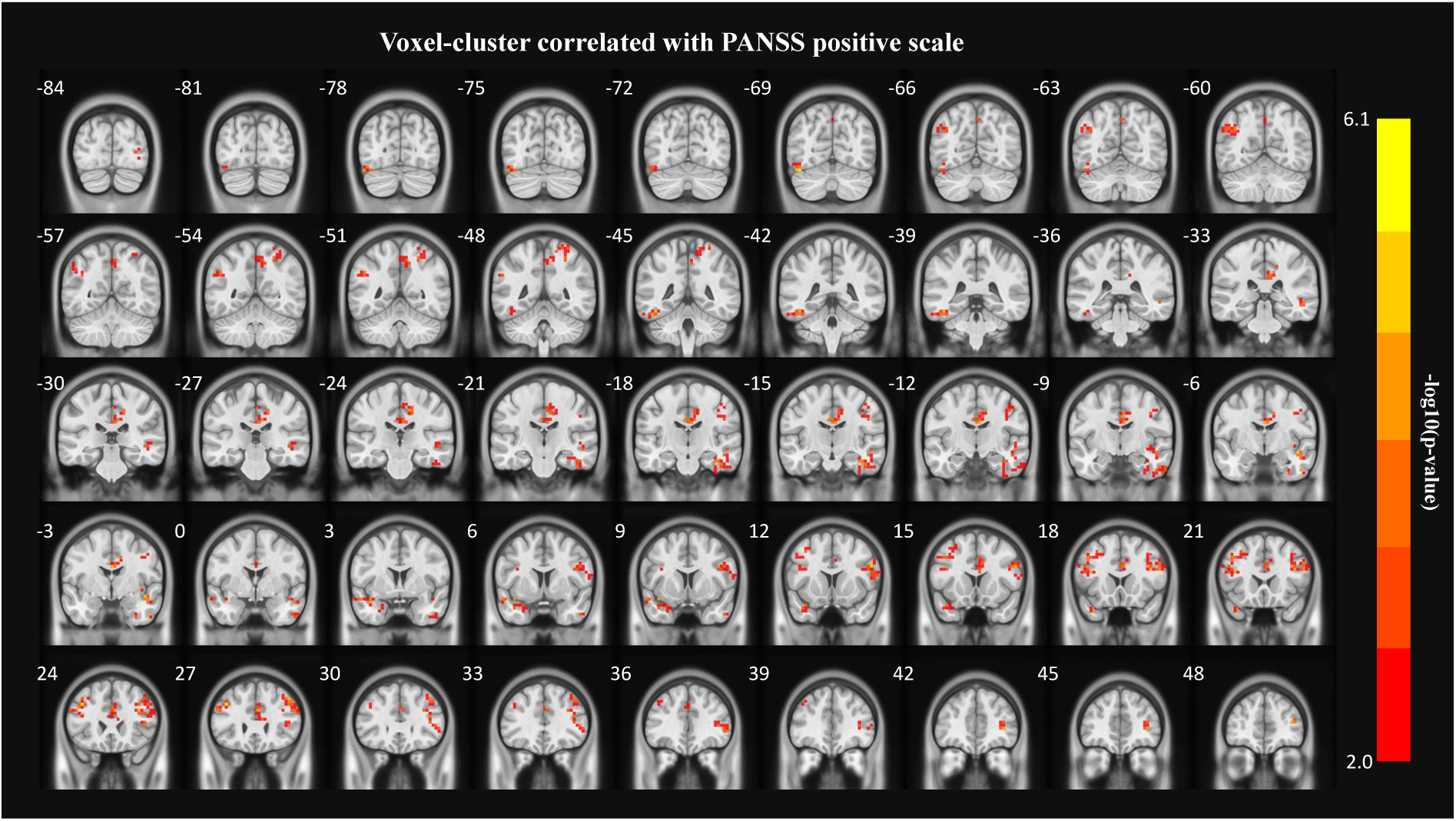
The voxel clusters with significant connectivity pattern correlations with the mean of the PANSS positive symptoms scores in schizophrenic patients across the COBRE and Taiwan datasets. The results were obtained using sKPCR controlling for the age, gender, mean FD and sites. The statistical maps were thresholded using permutation-based cluster-wise inference with a cluster-defining threshold of *p* = 0.01 and cluster size *p* < 0.05. The number to the left of each coronal slice is the MNI Y value.

**Figure D.15:**
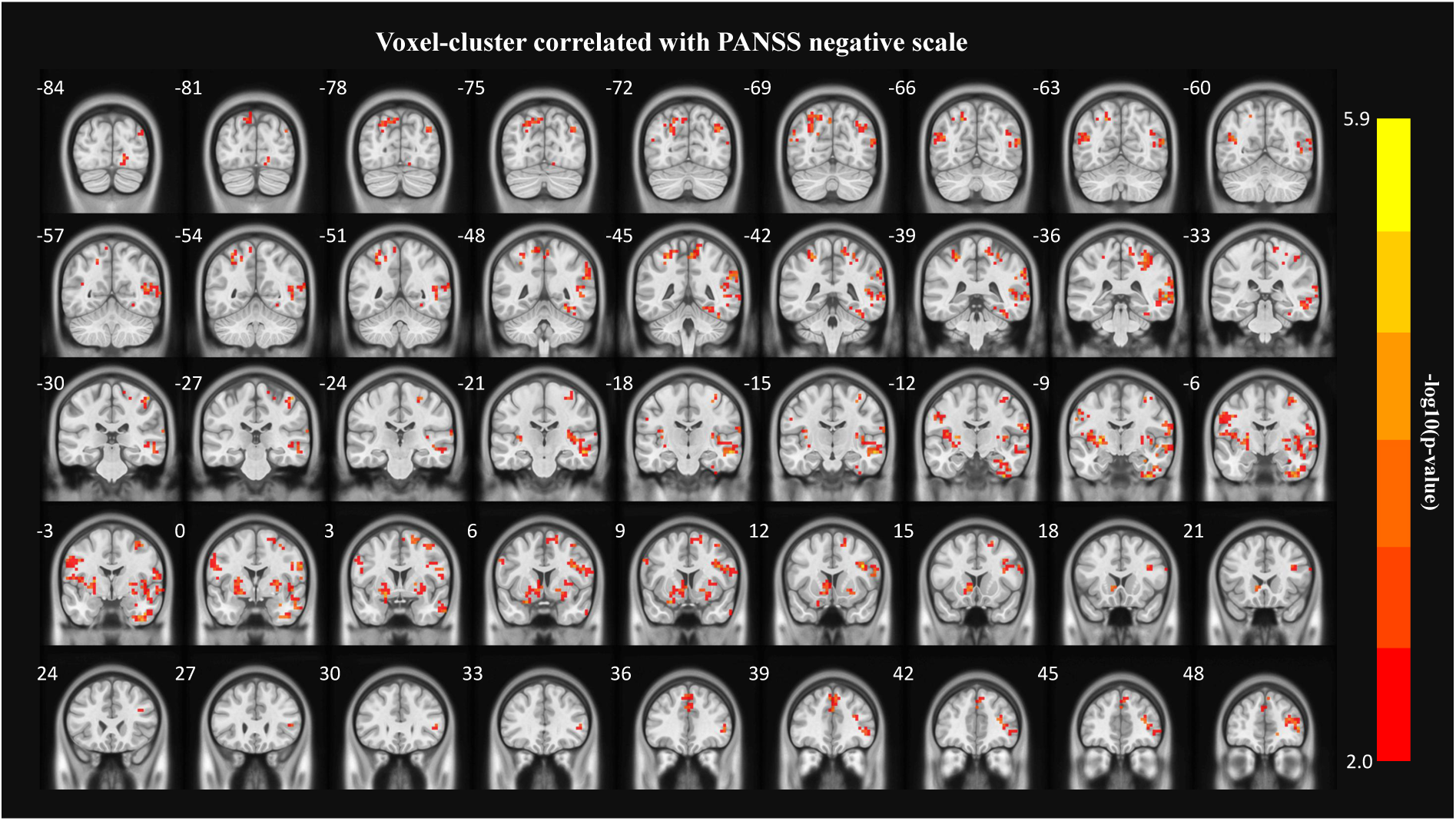
The voxel clusters with significant connectivity pattern correlations with mean of the PANSS negative symptom scores in schizophrenic patients across the two datasets. The results were obtained by sKPCR controlling for the age, gender, mean FD and sites. The statistical maps were thresholded using permutation-based cluster-wise inference with a cluster-defining threshold of *p* = 0.01 and cluster size *p* < 0.05.

## Footnotes

1 For computation time comparison, the aSPU is implemented in Matlab based on the function ‘aSPU’ in the R package ‘aSPU’.

